# MultiMatch: Geometry-Informed Colocalization in Multi-Color Super-Resolution Microscopy

**DOI:** 10.1101/2024.02.28.581557

**Authors:** Julia Naas, Giacomo Nies, Housen Li, Stefan Stoldt, Bernhard Schmitzer, Stefan Jakobs, Axel Munk

## Abstract

With recent advances in multi-color super-resolution light microscopy it has become possible to simultaneously visualize multiple subunits within complex biological structures at nanometer resolution. To optimally evaluate and interpret spatial proximity of stainings on such an image, colocalization analysis tools have to be able to integrate prior knowledge on the local geometry of the recorded biological complex. Here, we present *MultiMatch* to analyze the abundance and location of chain-like particle arrangements in multi-color microscopy based on multi-marginal optimal unbalanced transport methodology. Our object-based colocalization model statistically addresses the effect of incomplete labeling efficiencies enabling inference on existent, but not fully observable particle chains. We showcase that MultiMatch is able to consistently recover all existing chain structures in three-color STED images of DNA origami nanorulers and outperforms established geometry-uninformed triplet colocalization methods in this task in a simulation study. Furthermore, MultiMatch also excels in the evaluation of simulated four-color STED images and generalizations to even more color channels can be immediately derived from our analysis. MultiMatch is provided as a user-friendly Python package comprising intuitive colocalization visualizations and a computationally efficient network flow implementation.

## 1 Introduction

Colocalization analysis aims to unravel the interconnection and interaction network between two or more groups of particles based on their spatial proximity in a microscopy image. By visualizing biological structures, like DNA, RNA and proteins, that are only a few nanometers in size, colocalization analysis makes it possible to study a wide range of biological processes, such as DNA replication and the transcription of genes (Cainero et al, 2021), nuclear import of splicing factors (Costa et al, 2021) or the dynamics of cargo sorting zones in the trans-Golgi networks of plants (Shimizu et al, 2021), to name only a few.

In the following, we will denote any objects of interest that are depicted within a microscopy image, e.g., proteins as well as loci on DNA or RNA strands, as *particles*. In fluorescence light microscopy, such particles are stained, i.e., in case they do not already intrinsically fluoresce, they are labelled with fluorophores, which in turn are excited by an external light source. The emitted fluorescence radiation then can be imaged via several microscopy technologies.

Diffraction unlimited super-resolution fluorescence microscopy technologies, also called nanoscopy, are classified into two broad concepts (Sahl et al, 2017):

In **coordinate-stochastic microscopy**, fluorophores within the sample are stochastically excited resulting in a temporally resolved blinking dynamic (Betzig et al, 2006; Hess et al, 2006; Rust et al, 2006), which allows to spatially separate fluorophores. Their coordinates are estimated by means of the detected radiation peak, yielding a *list of coordinates of detected fluorophores* as output data. If only one fluorophore is detected for one particle, the output translates into a list of particle coordinates. Else, fluorophore coordinates can be aggregated in order to localize the particle of interest in the imaged biological sample.

In **scanning-based microscopy** methods such as Stimulated Emission Depletion (STED; Hell and Wichmann, 1994; Hell, 2007; Klar et al, 2000), the fluorescence distribution is stored as an *intensity matrix*, in which every entry encodes the detected radiation within a respective pixel of the microscopy image. To obtain coordinate estimates of particle positions, object detection algorithms have to be applied to the intensity matrix.

In order to study possible particle interactions or connections, stainings with different fluorescent markers are recorded in different color channels. Particles colocalize, if they are spatially closer than or equal to a *colocalization distance*, which heavily depends on the underlying biological setting and might be unknown prior to colocalization analysis (Malkusch et al, 2012).

Colocalization methods are divided in two categories based on the input data format they require:

**Pixel-based colocalization methods** take an intensity matrix as input and compare the pixel intensities across color channels, e.g., by utilizing overlap, correlation or intensity transport analysis. Such approaches are thus only applicable for scanning-based images and examples for well-established methods are Mander’s Colocalization Coefficient (Manders et al, 1993; Xu et al, 2016), Pearson’s Correlation Coefficient (Adler and Parmryd, 2010), BlobProb (Fletcher et al, 2010), SACA (Wang et al, 2019) and OTC curves (Tameling et al, 2021).

**Object-based colocalization methods**, which our method MultiMatch classifies as, require the coordinates of particles and evaluate their distances. Examples for other object-based tools are ConditionalColoc (Vega-Lugo et al, 2022) and Ripley’s K based methods (Ripley, 1976; Mukherjee et al, 2020) as SODA (Lagache et al, 2018).

Pairwise particle distances can be defined in several ways (Vega-Lugo et al, 2022). In MultiMatch we implemented the distance between reference points, i.e., the center of the detected particle, as default. However, we also allow the user to input a pre-defined particle-to-particle distance matrix, in case other approaches like the distance between object borders is preferred.

While nanoscopy for dual-color stainings is well studied for a long time, multi-color imaging including three or more stainings has received increased attention more recently since it allows simultaneous measurements of multiple particle types. There is a steadily increasing number of published multi-color STED microscopy datasets (Winter et al, 2017; Spahn et al, 2019; Butkevich et al, 2021; Glogger et al, 2022; Gonzalez Pisfil et al, 2022; Wang et al, 2022; Saal et al, 2023), of other super-resolution microscopy methods (Andronov et al, 2022; Unterauer et al, 2023) and the development of appropriate labeling methods allowing for an ever-increasing number of channels is ongoing (Beater et al, 2015; Butkevich et al, 2021; Willig et al, 2006; Reinhardt et al, 2023; Unterauer et al, 2023).

However, most pixel- and object-based colocalization tools are designed for and therefore limited to the analysis of two-color stainings. Applying them to multi-color images is not an obvious task: Particle arrangements with more than two different particle types can occur in different configurations (Figure 1B), and depending on the biological context, some may be of interest and others may simply not exist in the imaged sample. A geometry-uninformed, pairwise analysis of all possible channel combinations (Smallcombe, 2001), as well as the few established methods that are explicitly presented as multi-color pixel-based (Sastre et al, 2019; Goucher et al, 2005; Humpert et al, 2015; Fletcher et al, 2010) and object-based (Haas and Peau-celle, 2021; Lagache et al, 2018; Vega-Lugo et al, 2022) colocalization tools are prone to overestimate colocalization, as soon as the biological complex of interest has a fixed geometry and stoichiometry, as we can show in a simulation study (Figure 2A). To exploit the full potential of multi-color microscopy imaging in such a situation, it is therefore beneficial to actively incorporate prior knowledge of the local geometry into the colocalization analysis.

**Fig. 1.**
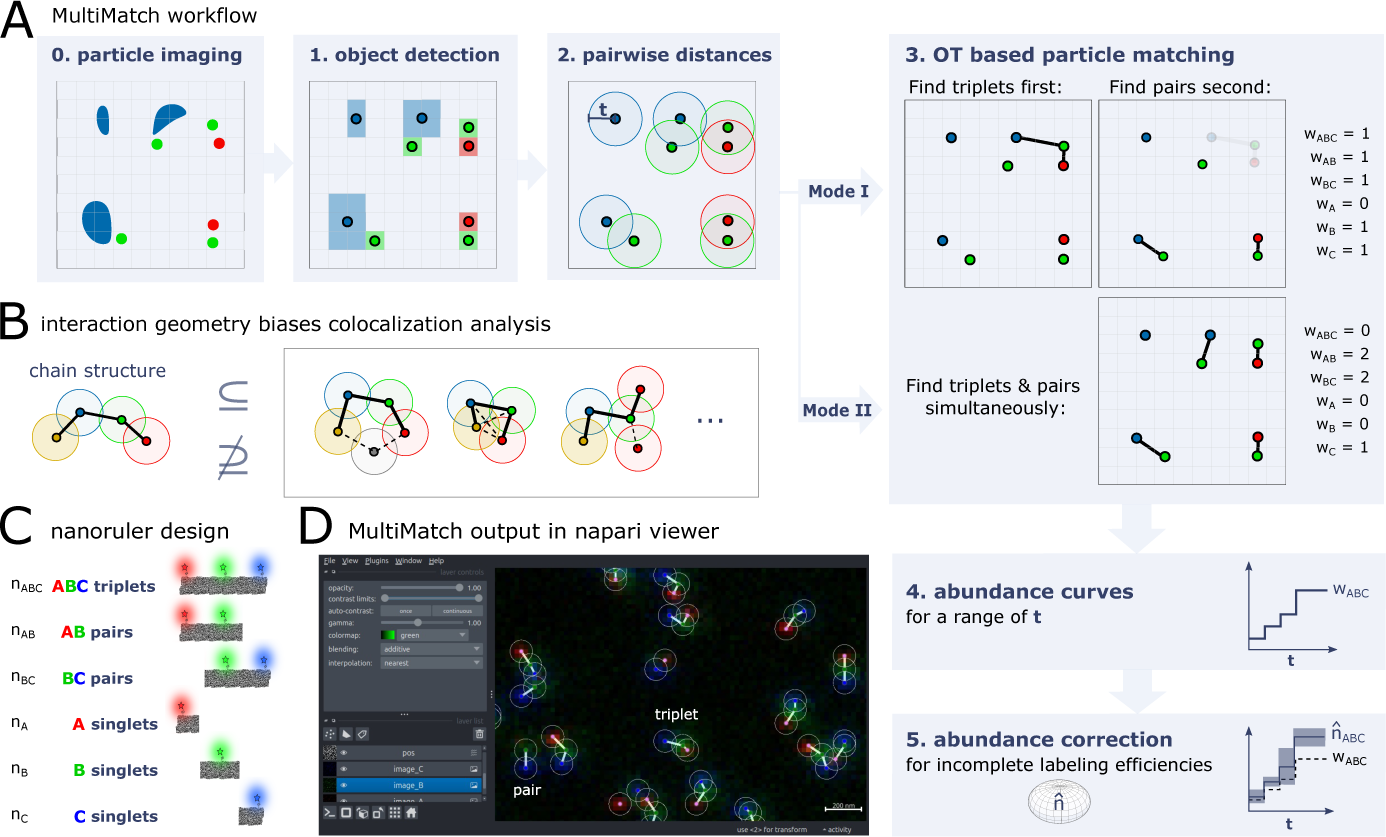
MultiMatch workflow to detect chain-like particle arrangements and experimental nanoruler design. **A.** After microscopy imaging (0) and object detection (1), the distances between channel-specific lists of reference points or a user-defined distance matrix are input to the optimal matching procedure. Restricted on particle pairs with distance smaller or equal than colocalization distance *t* (2), MultiMatch either outputs the maximal number of triplets and subsequently pairs (Mode I) or simultaneously searches for triplets and pairs (Mode II) (3). MultiMatch provides the localization and number of detected chains for a known or abundance curves for a range of colocalization distances *t* (4). For known incomplete labeling efficiencies true abundances can be estimated with confidence statements (5). **B.** If more than two different particle types are involved, multiple geometric colocalization patterns can emerge. In case the chain is a substructure of the colocalization geometry of interest, its detection will help to localize and quantify colocalization events. **C.** Structures of interest in three-color colocalization analysis for chain-like, one-to-one particle interactions and fixed particle type order. All pairwise distances between neighboring particles in a chain are smaller or equal than colocalization distance *t*. **D.** Exemplary MultiMatch output for an experimental STED image of DNA origami nanoruler structures (as sketched in C) in the interactive napari viewer.

To this end, we introduce MultiMatch, a widely applicable colocalization methodology based on optimal transport theory, which is especially tailored to detect chain-like, one-to-one particle arrangements. Integrating this type of colocalization geometry optimizes the multi-color colocalization analysis of quadruples, triplets, pairs, and singlets, as they appear when marking different loci of a chain-like molecule with multi-color stainings.

One exemplary biological framework, in which the localization of such arrangements is especially insightful, is the highly condensed mammalian mitochondrial genome: It is transcribed from both strands of the mitochondrial DNA as long polycistronic transcripts that have to undergo multiple processing steps, including endonucleolytic cleavage, in order to get to the different functional RNA species. Transcription of the heavy strand leads to polycistronic primary transcripts containing the premature mRNAs of 12 of the 13 OXPHOS subunits encoded in the mitochondrial genome. Labeling more than two of the mRNAs within such a primary construct, in combination with our novel colocalization approach, can significantly contribute to our understanding of the post-transcriptional processing steps and their dynamics, that lead to the generation of matured mRNA molecules (Boettiger et al, 2016; Miron et al, 2020).

However, even if the biological complex of interest itself is not chain-like, chain detection still can give substantial insights on the abundance and location of colocalization events inside a microscopy image as soon as the chain is a substructure of the colocalization geometry (Figure 1B). The converse, on the other hand, does not hold true in general.

We consider a particle arrangement as *chain-like* when exactly one particle of each type is stringed together in an ordered fashion and pairwise distances of chain-neighbors are smaller than or equal to a maximal colocalization threshold *t*. To fix the chain order of particles, we will refer to color channels, in which the respective particle type was imaged, as channel A, B, C, D etc.. For simplicity, we will explain the main methodology for a three-color setting in what follows, but MultiMatch is applicable to an arbitrary number of color channels, which we showcase in the evaluation of simulated four-color STED images (Section 2.5). We stress, that our software (see Section 4.8) is already designed to process any number of channels.

All configurations resulting from a three-color staining of an chain-like molecule are sketched in Figure 1C, where we assume the following unknown abundances ***n*** = (*n_ABC_, n_AB_, n_BC_, n_A_, n_B_, n_C_*) of chain-like assemblies, where

*n_ABC_* is the number of true ABC triplets,
*n_AB_, n_BC_* is the numbers of true AB and BC pairs,
*n_A_, n_B_, n_C_* is the numbers of true A, B and C singlets.

MultiMatch outputs detected abundances ***w*** = (*w_ABC_, w_AB_, w_BC_, w_A_, w_B_, w_C_*) for a known colocalization distance *t* and depicts configuration positions on the respective microscopy image allowing further investigation on the spatial distribution of recorded biological com-plexes. If *t*is unknown (optionally channel-wise scaled) abundance curves ***w***(*t*) are output for a user-defined range of *t* values. MultiMatch is compatible with the interactive Graphical User Interface of napari (Figure 1D) enabling the visual evaluation of structure locations for different *t* values in form of a colocalization threshold slider.

The differentiation between triplets, pairs, and singlets within a microscopy image is additionally hindered by incomplete labeling efficiencies and point detection artifacts. This is a notorious problem in fluorescence microscopy, e.g., described in Hummert et al (2021), and missing detections can add an unpredictable bias towards systematic underestimation of triplet numbers and overestimation of singlet abundances, if not corrected. Currently, the problem of incomplete labeling efficiency is barely addressed in the field of colocalization analysis.

Therefore, we propose a statistical framework to correct for incomplete labeling efficiencies and introduce an unbiased estimator ***n̂***(*t*) of true chain-structure abundances and confidence statements on the estimated quantities. An overview on the full workflow of MultiMatch from microscopy image to abundance curves is depicted in Figure 1A.

## 2 Results

### 2.1 Chain-like Particle Assembly Detection with MultiMatch

Optimal transport (OT) theory (Villani, 2009) has a wide range of applications throughout statistics (Panaretos and Zemel, 2019), data science and machine learning (Peyŕe and Cuturi, 2020). Generally, OT aims to allocate (transport plan) one mass distribution into another by minimizing the transportation cost arising from moving one mass unit from one location to another. Applied to fluorescence intensity distributions on a pixel grid and using the euclidean distances between pixels as transportation cost, OT introduces an intuitive distance between two microscopy images and could already successfully be utilized in the context of pixel-based, dual-color colocalization methods (Zaritsky et al, 2017; Tameling et al, 2021).

For object-based analysis, reference points of detected particles can also be interpreted as support points of mass one of a (discrete) two-dimensional distribution. For only two color channels with the same number of particles the standard OT problem simply assigns each particle from the first channel to one particle from the second channel while minimizing the total sum of Euclidean matching costs. We can obtain an optimal matching between more than two particle types by multi-marginal OT (Kim and Pass, 2014; Pass, 2015) and at the same time account for the not necessarily equal numbers of support points per channel by utilizing an unbalanced OT formulation (Chizat et al, 2018). A combination of both OT generalizations, i.e., multi-marginal optimal unbalanced transport problems, have been recently discussed in the literature (Friesecke et al, 2021; Heinemann et al, 2022; Beier et al, 2022; Le et al, 2022).

In this manner, the basic concept of MultiMatch can be interpreted as linear assignment problem as described, e.g., in the field of object tracking (Schulter et al, 2017; Chari et al, 2015; Jaqaman et al, 2008; Zhang et al, 2008). In contrast to methods of this research field, we explicitly formulate the matching problem as a function of the colocalization threshold, allowing to plot the chain abundances dependent on a range of *t*. Furthermore, we develop a novel statistical framework specific to labelled marker colocalization to infer on the statistical influence of incomplete labeling efficiency. We utilize the equivalence of the optimal transport methodology to a network flow problem to overcome the otherwise prohibitively high computational complexity of its corresponding linear program formulation (Lin et al, 2022; Supplementary Material A.2).

MultiMatch provides two different modes to solve the particle matching problem (Figure 1A(3)):

**Mode I:** By restricting a *k* -marginal optimal unbalanced transport problem to particle pairs with a distance smaller than *t* and introducing a chain-cost that only considers distances between neighboring particle types (Supplementary Material A.1), the resulting OT plan encodes the *maximal* number of, for *k* = 3, triplets within the nanoscopy image. If requested, the matching process is subsequently repeated on the remaining particles to detect yet unresolved AB and BC pairs, respectively.

**Mode II:** This mode only detects AB, BC, etc. pairs by solving respective *two*-marginal unbalanced OT problems. Subsequently, the two-marginal OT matchings are coupled to chain structures: For *k* = 3, all pairs occupying the same intermediate particle are redefined as respective ABC triplet.

Depending on the underlying biological experiment, the user can select the appropriate mode for colocalization analysis: Mode I prioritizes the detection of a predefined chain structure of choice. For example, if a user aims to analyze triplets, Mode I will detect a triplet as soon as three particles A, B, and C are close enough to each other – even if another particle A or C is nearby that would allow to match two pairs instead of one triplet (as depicted in Figure 1A(3)). If *k >* 3 and the user wants to detect multiple chain structures, one needs to set a prioritization order for Mode I. For example, for *k* = 4 and after ABCD quadruplet detection, one can search either for ABC or BCD triplets next. Depending on the order, the final matching results may change as soon as some particles cannot be uniquely assigned to one particle arrangement.

Mode II, on the other hand, does not need a predefined prioritization order of structures for subsequent matching steps, hence it does not overemphasize structures that are matched in the earlier steps. It is useful in case we do not have any prior knowledge on which structures might appear in the microscopy image and we do not want to prioritize any chain structures.

In the evaluation of experimental (Section 2.4 and Supplementary Material D, Figure D4) and simulated three-color STED microscopy images (Figure 2 and Supplementary Material E.1, Fig. E5) we show that for sparse particle distributions and mixed singlets, pairs, and triplet ratios the differences in detected abundances between the two modes is neglectable. However, in case of dense particle distributions (see Supplementary Material E.2, Fig. E6 and Supplementary Material E.3, Fig. E7 B-D), or in case we know in advance that only one chain structure exists in the biological context, the multi-marginal approach of Mode I, which is also the default setting in the MultiMatch tool, outperforms the pairwise matching approach of Mode II.

**Fig. 2.**
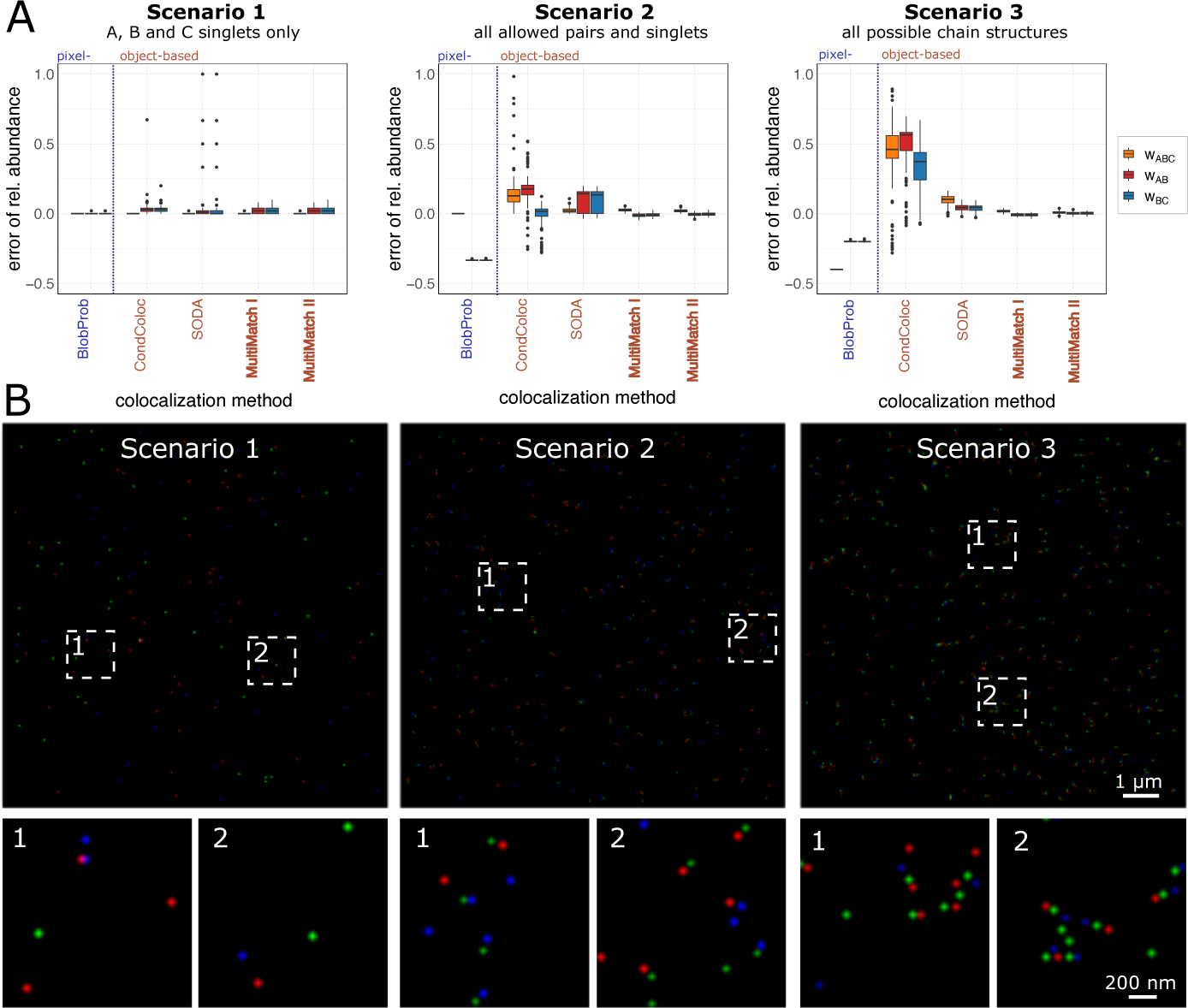
Simulation study for three combinations of chain structures. In each Scenario 100 STED images and different abundances of triplets, pairs, and singlets were sim-ulated with 100% labeling efficiency. **A.** Method specific boxplots of the errors in detected relative (scaled by the total number of points in channel B) structure abundances are dis-played. The error is computed by subtracting true relative abundance from detected relative abundances. In *Scenario 1* only A,B and C singlets, in *Scenario 2* all possible singlets as well as AB and BC pairs and in *Scenario 3* ABC triplets, AB, BC pairs and A, B and C singlets were simulated. **B.** Simulated STED images for Scenarios 1, 2 and 3 with respective image details.

### 2.2 Simulation Study

To systematically evaluate the performance of MultiMatch against compatible colocalization methods, we simulated 100 microscopy images for each of three scenarios with different combinations of singlets, pairs, and triplet abundances. For this simulation study, we decreased the noise level to a minimum to allow a fair comparison despite different point detection tools implemented in the respective colocalization tools. Also, we amplified simulating linear triplet structures over randomly folded triplets (see simulation setup in Section 4.4). For every simulated image,

**Scenario 1:** 50 singlets of each type A, B and C were simulated.
**Scenario 2:** 50 A, B and C singlets and 50 AB and BC pairs were simulated, respectively.
**Scenario 3:** 100 triplets and 50 AB and BC pairs and 50 A, B and C singlets were simulated, respectively.

Exemplary, simulated images and the results of the simulation study for a fixed colocalization threshold of *t* = 5 pixels are shown in Figure 2. Analysis results for all considered methods across a range of colocalization thresholds are presented in Supplementary Material E.1, Figure E5.

As a representative of pixel-based methods, we include BlobProb (Fletcher et al, 2010), which counts the number of colocalized intensity blobs, i.e., groups of neighboring pixels with high intensity. In each channel, blobs are detected via image segmentation and for each blob the local intensity maximum is defined as reference particle coordinate. A blob pair colocalizes if the first reference point lies within the second blob and vice versa. Triplet colocalization is detected if all involved reference points are included in all three blobs. SODA (Lagache et al, 2018) is an object-based method, which uses the Ripley’s K function (Ripley, 1976) and computes the coupling probability of point pairs based on marked-point process theory. In the most recently published method ConditionalColoc (Vega-Lugo et al, 2022) particles are defined as colocalized as soon as their distance is below a maximal colocalization radius. Then, utilizing Bayes’ Theorem, (conditional) probabilities are computed and assigned for triplet and pair colocalization. We experienced that ConditionalColoc, although aiming to output probabilities, in some cases yields values greater than one and hence the errors in relative abundance detection are not bounded by one as well. For a better comparison, we restricted the respective results to values between −0.5 and 1 in Figure 2 and show ConditionalColoc outliers in Supplementary Material C (Figure C3).

In none of the above methods triplet colocalization is restricted to one-to-one interactions. This has barely any negative effect on the detection of singlets in Scenario 1, where no additional pairs and triplets occur. Apart from few outliers of overestimation in pairs and triplet abundances in ConditionalColoc and SODA, all considered colocalization measures show consistently low errors with small variability. The maximal median error in relative abundances of 0.03 in Scenario 1 is obtained by ConditionalColoc in the detection of AB as well as BC pairs.

In Scenarios 2 and 3 on the other hand, we observe a consistent overestimation of relative pairs and triplet abundances in object-based methods SODA and ConditionalColoc, since one particle can be included in several structures at the same time. Additionally, in Scenario 2 SODA exhibits a larger variation in pairs abundances, resulting in median errors 0.14 in both AB and BC pairs with interquartile ranges of 0.16, respectively. In Scenario 3 the variation in abundance detection decreased and median errors are 0.1 for ABC triplets and 0.04 for AB as well as BC pairs. ConditionalColoc performances worst in Scenario 3 yielding a median error of 0.48 for ABC triplets.

The pixel-based method BlobProb mostly obtains zero relative abundances of triplets and pairs across all three scenarios and hence severely underestimates the triplet and pair configurations within the simulated images. This is due to the high resolution in the simulation setup, which was chosen to mimic conventional STED imaging. If particles are small and their respective intensity blobs do not overlap, BlobProb does not detect any colocalization.

MultiMatch on the other hand searches for optimal matches on a global scale while considering the local geometry of chain-like particle assemblies. It consistently recovers the ground truth abundances for each simulation scenario. The maximal median error across all scenarios and chain structures for both Modes of MultiMatch is 0.03 with a maximal interquartile range in errors of 0.04.

Apart from above considered, already established colocalization methods, we also implemented a Nearest Neighbor Matching as comparable object-based method. We can show that greedily matching particle pairs based on local optima leads to underestimation of ABC triplets in dense particle distributions (Supplementary Material E.2, Figure E6A-C).

### 2.3 Incomplete Labeling Efficiencies and Point Detection Errors

In experimental STED microscopy, typically it is impossible to record all exist-ing particles of interest. This can, for example, be due to the fluorescent marker not being successfully attached to the probe or a flawed point detection. All such scenarios resulting in a failure of particle detection for simplicity will be summarized under *incomplete labeling efficiency* hereafter.

If only singlets were to be counted in multi-color images with the same labeling efficiency across channels, the relative abundance could still be estimated consistently. However, as soon as configurations of two or more particle types are to be recovered, incomplete labeling efficiencies can lead to under- and overestimation of structures. Figure 3A shows that a triplet can be erroneously detected as pair or singlet or not at all, which can introduce a severe bias. However, if the labeling efficiencies are known, the detection success of a particle can be modeled with a Bernoulli distribution, which allows the definition of an unbiased estimator ***n̂*** for the vector of true chain structures abundances ***n***. This approach allows for constructing multi-dimensional joint confidence ellipsoids covering ***n*** with a given significance level, e.g., *α* = 0.1 (Figure 3B,C). The multi-dimensional confidence ellipsoid then can be respectively projected onto one dimension to obtain structure-specific confidence intervals or bands for a range of *t* values, while fixing the estimated abundances of all other considered structures. For more details on the statistical framework see Supplementary Material B.

**Fig. 3.**
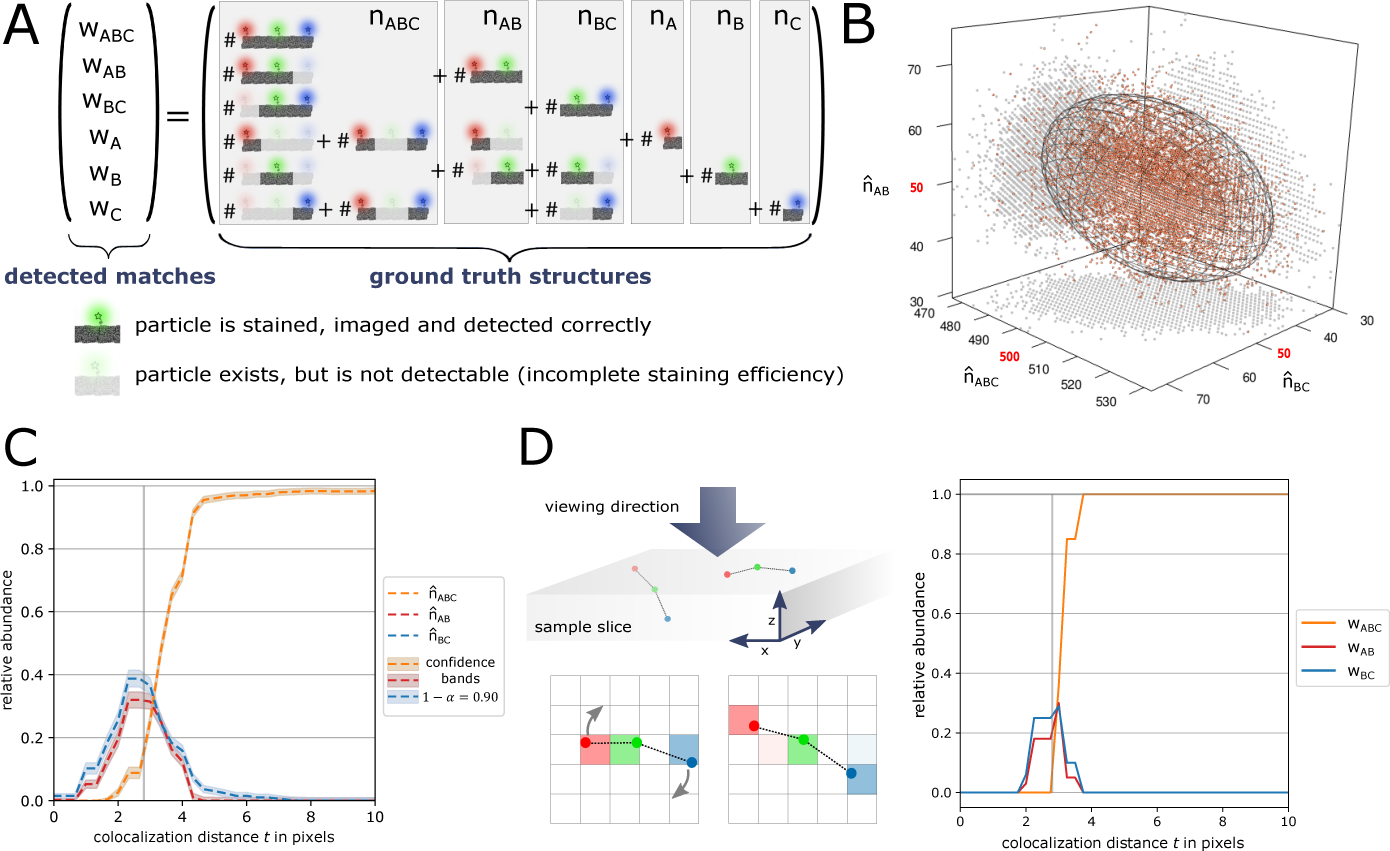
Incorporating incomplete labeling efficiency. **A.** Because of channel-specific incomplete labeling efficiencies, triplets and pairs can erroneously counted to other structure abundances. **B.** For entrywise large enough ***n***, estimator ***n̂*** is approximately multi-dimensional normally distributed: Estimated abundances of 10,000 simulations with labeling efficiencies *sA* = *sB* = *sC* = 0.95 and true abundances *nABC* = 500*, nAB* = *nBC* = *nA* = *nB* = *nC* = 50 (see Supplementary Material B.2). The respective 3-dimensional, normal 90% quantile ellipsoid is plotted. **C.** Estimated abundance curves for one of the experimental multi-color STED images in Setting 3 with additional confidence bands for significance level *α* = 0.1. **D.** Restricted image resolution and 3-dimensional rotation of particle arrangements lead to variability in the observed colocalization thresholds: Simulation study of 100 images only containing one triplet with pairwise distances set to 70 nm = 2.8 pixels per image (100% complete labeling efficiency, see Section 4.4).

### 2.4 Evaluation of Experimental STED Images

Chain-like particle structures occur within several biological complexes. To showcase the performance of our method on experimentally retrieved data we used one-, two- and three-color nanorulers. Nanorulers are DNA-origamis with a predefined distance between spots at which 20 fluorophores are attached and hence, as their name suggests, can be used as rulers inside a microscopy image (Cainero et al, 2021; Schmied et al, 2014, 2012; Rothemund, 2006). For this experimental setup, we chose nanorulers with pairwise distances between neighboring spots of 70 nanometers (nm). For each chain structure (as depicted in Figure 1C), respective nanoruler origamis are available in separate solutions, which allows us to control whether in an experiment we record singlets, pairs or triplets only or a combination of those structures. We performed three experiments:

**Setting 1:** The experiment consists of all three single marker nanorulers (22 images in total). We expect to detect no pairs or triplets, i.e., *w_ABC_* = *w_AB_*= *w_BC_* = 0.
**Setting 2:** The experiment consists of all three singlets, two pairs and triplet marker nanoruler solutions (22 images in total). We expect to detect all possible configurations, i.e., A, B and C singlets, AB and BC pairs as well as ABC triplets.
**Setting 3:** The experiment consists of only triplet marker nanorulers (12 images in total). We expect to detect ABC triplets only, i.e., *w_AB_* = *w_AB_* = *w_A_*= *w_B_* = *w_C_* = 0.

For each experimental setting we recorded STED images of size 400 *×* 400 pixels with a pixel size of 25 *×* 25 nm. In channel A, stainings with Star Red 640 nm are recorded, in channel B, stainings with Alexa 488 and in channel C, stainings with Alexa 594. Note, however, that the exact numbers of nanorulers within a recorded STED image is unknown. Due to misfolding and clumping of nanorulers and different nanoruler immobilization rates for each STED image one cannot compute a fixed unit of nanorulers per microscopy image and experiment.

The results of the colocalization analysis for all three settings (with default MultiMatch Mode I) are shown in Figure 4 via relative abundance curves with standard deviation bands quantifying variation across images within the same setting. Here, we used MultiMatch Mode I and included the analysis with Mode II showing comparable results, but slightly underestimating the number of triplets in Setting 3, in Supplementary Material D, Figure D4.

**Fig. 4.**
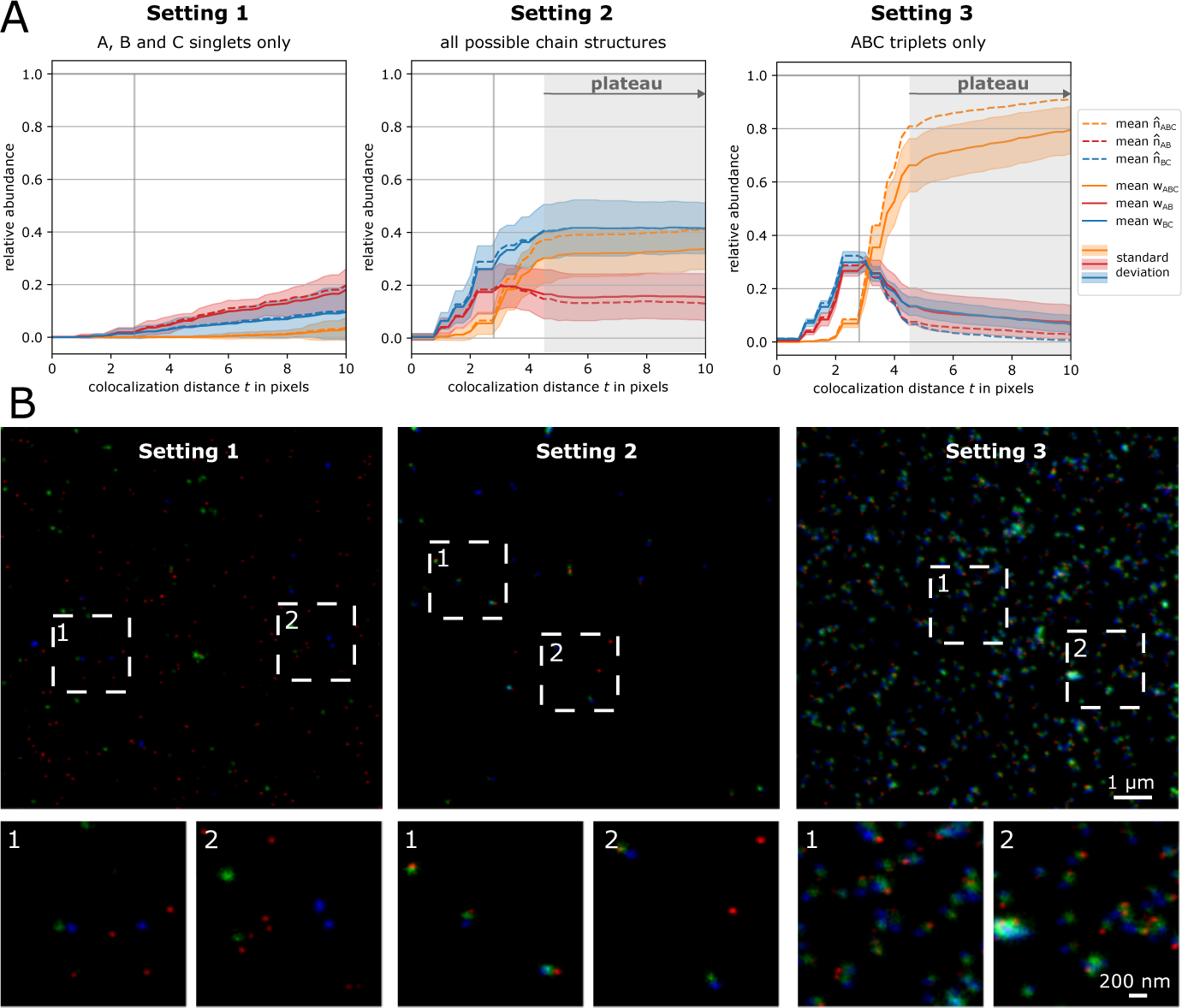
MultiMatch Mode I relative abundance curves *w*(*t*) for experimental STED images. For each setting the solid curves are mean relative abundances with standard deviation bands across a range of colocalization threshold *t* from 0 to 10 pixels (25 nm = 1pixel). The abundances are scaled by the total number of points detected in channel B. Additionally, incomplete labeling efficiency (90% in each channel) corrected abundances are plotted as dotted curves. The true colocalization distance of 70 nm within nanoruler structures is depicted as vertical line. **A.** *Setting 1:* Mean abundance curves for only singlets consistently show the expected 0% relative triplet and pair abundances. *Setting 2:* Triplets, pairs, and singlet nanoruler are detected with stable abundances for approximately *t ≥* 4 pixels. *Setting 3:* Mean abundance curves for analyzing the triplet nanoruler solution only. The incorporation of incomplete labeling efficiency clearly corrects the relative triplet abundance towards the in this setup expected 100%. **B.** Representative STED images for Settings 1,2 and 3 with image details.

For Setting 1 we can appreciate that, as expected, across a range of *t* values only a few pairs and triplets are detected (Figure 4A). The rise of relative abundance curves is unavoidable for large *t*, since the probability increases that randomly scattered particles are matched. In Setting 2, despite experimental variation, we clearly recover all supplied nanoruler structures. Even more, colocalization curves are still stabilizing for a colocalization threshold *t* greater than approximately 4 pixels (= 100 nm): For *t >* 100 nm ABC triplets are approximately detected with relative abundance of 0.32, AB pairs with 0.16 and BC pairs with 0.42 relative abundance, yielding a relative amount of 0.1 unmatched B singlets. The relative abundance curves of all structures reach a plateau at approximately *t ≥* 4 pixels (= 100 nm), i.e., the slope of all curves within the same setting decreases rapidly. In Setting 3, as expected, the relative abundances of AB and BC pairs converge to zero while triplets are the dominantly detected structure for *t ≥* 4 pixels.

Notably, in Settings 2 and 3 stable abundance curves are reached at around 100 nm, which is 30 nm more than the experimentally fixed, maximal distance between neighboring fluorophore spots in the nanoruler structures. This effect can be explained by the still limited resolution in the microscopy image and can be reproduced via simulation: We simulated 100 STED images containing only one triplet (*n_ABC_* = 1) in Figure 3D and can reproduce this stabilizing behavior of abundance curves in Figure 2.

Limited resolution alone does not explain why 20%–30% of detected B particles (for *t ≥* 5 pixels) are not matched to a triplet in Setting 3: The attachment of a single fluorophore to a nanoruler spot is expected to have a success probability of 85% to 90% and hence at least one fluorophore should be attached to each spot in almost 100% of all cases. Still, due to the above described experimental variation in nanoruler imaging and additional errors in point detection, especially due to nanoruler clumping, the overall success rate of fluorophore spot detection is incomplete. Hence, we erroneously detect pairs instead of triplets or singlets due to noise. As in Setting 1 those artifacts will be matched into triplets for large enough *t*.

For simplicity, we model a 90% labeling efficiency across all three-color channels in the experimental STED setup. The estimated abundance curves ***n*^**(*t*) (dotted lines in Figure 4), in Setting 3 visibly correct the measurements towards the expected relative abundances. Additional confidence bands around ***n*^** allow to infer on the robustness of the abundance estimation as presented in (Figure 3C) for one of the experimental STED images of Setting 3.

### 2.5 Evaluation of Simulated Four-Color STED Images

MultiMatch is applicable to an arbitrary number of color channels, which we showcase in the following with an adapted simulation setup for quadruples, triplets, pairs, and singlets in simulated four-color STED microscopy images. In contrast to the simulation study in Section 2.2, we additionally challenged our MultiMatch tool with an increased noise level and by allowing arbitrarily curled chain structures (see simulation setup in Section 4.4). In Figure 5 we show the colocalization analysis results of two simulation scenarios:

**Fig. 5.**
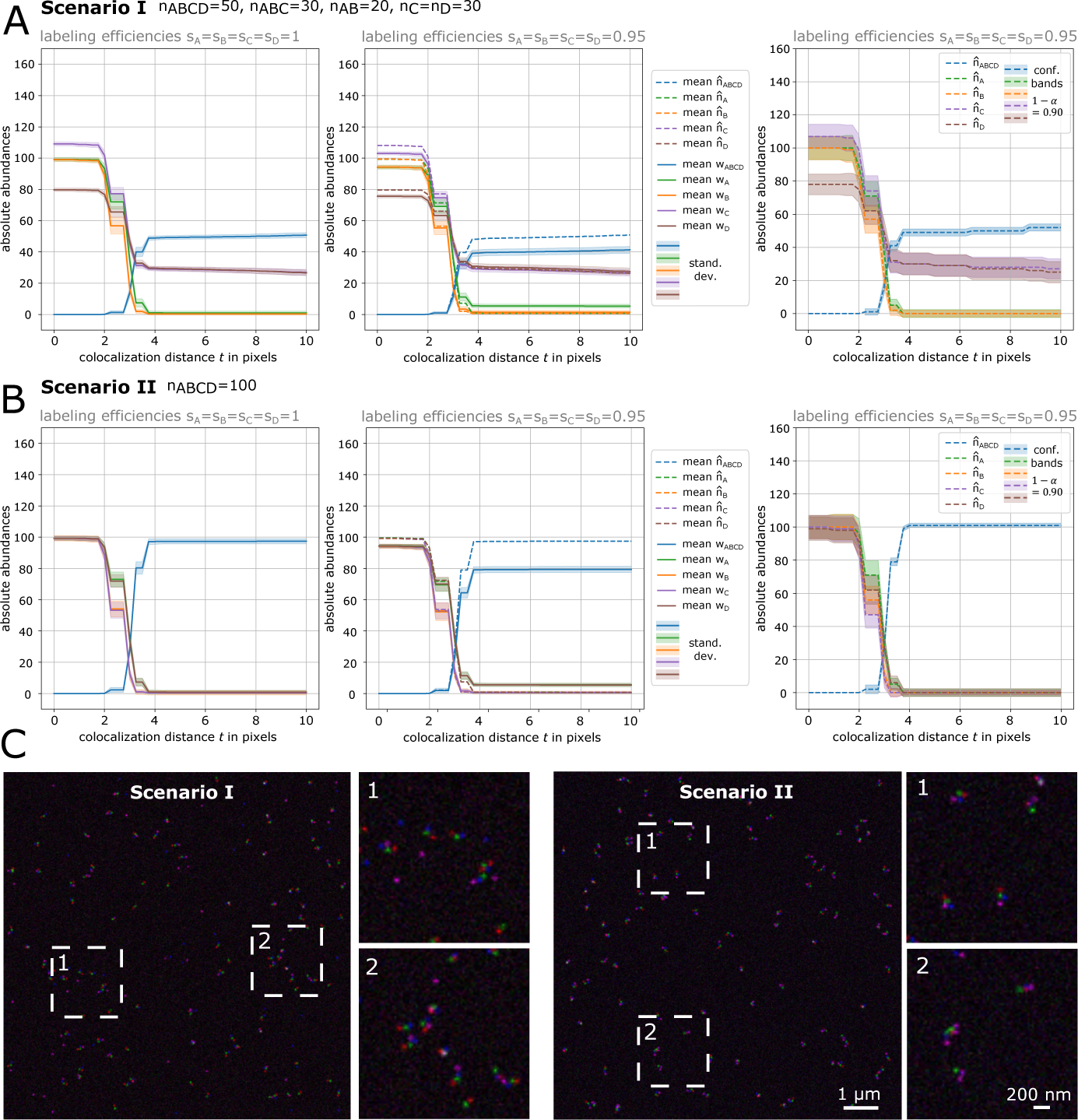
MultiMatch Mode II abundance curves *w*(*t*) and estimation results *n*^(*t*) for simulated four-colour STED images. *Scenario I:* A mixture of ABCD quadruplets, ABC triplets, AB pairs and C,D singlets were simulated. *Scenario II:* Only ABCD quadruplets were simulated. **A.** For each scenario images with complete labeling efficiency (left) and with incomplete labeling efficiency (middle) were simulated. Solid curves are mean absolute detected abundances with standard deviation bands across a range of colocalization thresholds *t* from 0 to 10 pixels (25 nm = 1pixel). All curves stabilize at approximately *t* = 4 pixel close to the true simulated number of structures. For images with incomplete labeling efficiency (95% in each channel) uncorrected detected abundances plus standard deviation bands are plotted as solid curves showing consistent underestimation of quadruples. Corrected abundances are plotted as dotted curves recovering the true number of simulated structures. For one exemplary STED image simulated with incomplete labeling efficiency, corrected abundance curves and corresponding confidence bands are shown (right). **B.** Representative STED images for Scenarios I and Scenario II with image details.

**Scenario I:** We simulated 50 ABCD quadruples, 30 ABC triplets, 20 AB pairs and 30 C and D singlets, respectively, to mimic a chain-like molecule being split at loci C and D.
**Scenario II:** We simulated 100 ABCD quadruples and no triplets, pairs nor singlets

Three additional simulations setups are shown in Supplementary Material E.3 and analysis results are plotted in Figure E7. For each scenario we simu-lated 100 images with full labeling efficiencies (*s_A_* = *s_B_*= *s_C_* = *s_D_* = 1) and 100 images with incomplete labeling efficiencies (*s_A_* = *s_B_*= *s_C_* = *s_D_* = 0.95) by randomly deleting 5% of points simulated in the prior, full labeling efficiency simulation in each channel.

For this analysis we applied MultiMatch Mode II, i.e., allowing the detection of both ABC as well as BCD triplets and AB, BC and CD pairs without any prioritization order of chain structures. Again, also in the case of four-color images, we can appreciate that MultiMatch consistently recovers true abundances of quadruplets in case of full labeling efficiencies. Absolute abundance curves, as also described in the analysis of our experimental dataset in Figure 4, stabilize for approximately *t* = 4 pixels. For images simulated with incomplete labeling efficiencies, the colocalization curves show underestimation of quadruplets as expected. With our statistical framework we again can visibly correct the colocalization curves towards the true, simulated structures abundances and additionally gain confidence bands confirming the stability of our estimator.

For denser distributions, as shown in Supplementary Material E.3, Fig.E7 B-D, we can observe that 1. MultiMatch II misses quadtruples for the sake of closer particle pairs, and 2. similar to the experimental nanoruler anaylsis depends on the performance of the point detection and hence the noise level in the microscopy image. If consistent noise challenges the point detection, abundance curves still stabilize, but the plateau shows a smaller number of matched quadtruples than simulated in absolute numbers. Hence, we advise user of MultiMatch to check the noise level of the microscopy image and the point detection result with the interactive napari viewer (Figure 1D and Supplementary Material E.3, Fig E7 D) and if necessary evaluate channel-wise scaled, relative instead of absolute abundances.

## 3 Discussion

In this article we introduce multi-marginal optimal unbalanced transport methodology for geometry-informed, multi-color colocalization analysis. We are able to show, that for the analysis of more than two color channels, it is crucial to take into account the colocalization geometry of the biological complex.

By either choosing chain-costs in a multi-marginal OT problem (Mode I) or coupling consecutive two-marginal OT matchings (Mode II), MultiMatch successfully detects *k*-chain particle assemblies such as quadruples, triplets, pairs, and singlets, as they appear when staining multiple loci on chain-like molecules like DNA or RNA strands. Both modes have their advantages, which depend on the number of particles imaged and prior knowledge on the biological context: Mode I is best for detecting one chain structure of choice and is more robust in dense particle distributions. When the particle distribution is sparser and multiple chain structures in the imaged biological setting are of interest, Mode II is suited to detect them without any predefined prioritization order.

Since often the true colocalization distance is unknown, MultiMatch results can be output as structure-wise relative or absolute abundance curves across a range of colocalization thresholds *t*. In our simulation studies as well as our experimental settings we could show, that output curves stabilize close to ground truth abundances.

However, as for all object-based colocalization methods, the performance MultiMatch scales with the noise level of the microscopy image, the performance of the object detection and the resolution of the microscopy. Abundance curve plateaus can be less clear in case the microscopy image contains detected singlets of different particles types. In this case, the larger *t*, the more far away singlets are matched. In such cases it might be unclear, whether singlets truly exist in the biological sample or whether they are an artifact of the experiment and image processing. For such cases, we advise to observe the quality of the microscopy image with the MultiMatch compatibale, interactive napari viewer.

Our network flow implementation significantly decreases computational costs compared to standard approaches solving comparable OT problems and comparable colocalization tools (Section 4.3, Supplementary Material A.2 and E.1). The simulation studies show that as soon as we have prior knowledge on the chain colocalization geometry, MultiMatch, in contrast to other triplet colocalization methods, is robust against overestimation of triplets with chain geometry since it only considers one-to-one interactions. MultiMatch is also tested on experimental STED images of different nanoruler combinations and can correct structure abundances for predefined incomplete labeling efficiencies and point detection errors, where confidence bands allow further inference on the estimated abundances.

All experimental studies have been performed for *k* = 3 color channels. However, in many scientific fields the detection of *k*-chains for larger *k* is of interest. The mathematical and statistical frameworks allow straight-forward generalization (Details in Supplementary Material A.1) and we exemplarily show successful detection results for simulated four-color STED images. With current technical standards, the experimental setup of multi-color nanoscopy imaging is still challenging, costly and time consuming, but in view of further technological improvements our algorithm is already applicable for the evaluation of this type of experimental setups, and especially promising in view of recent developments in super-resolution microscopy with a resolution of a few nanometers and below (Balzarotti et al, 2017; Gwosch et al, 2020).

In the same way channel specific colocalization thresholds as *t_AB_, t_BC_* and *t_CD_* can be considered within the OT problem. Although we only present the evaluation of 2D STED images with constant labeling efficiencies across channels, our software package can directly be applied to multi-color 3D microscopy images with channel-specific labeling efficiencies.

Limitations: If the microscopy image shows especially dense point clouds, MultiMatch necessarily will have difficulties in differentiating between random and biological reasonable proximity. Note, however, that this is not a specific weakness of MultiMatch, but any other method will face this identifiability problem, which is caused by missing linkage information. It can only be overcome with additional prior information of the underlying biological sample. However, MultiMatch Mode I is especially robust against dense particle distribution in comparison to pairwise matching approaches as implemented in MultiMatch Mode II or greedy Nearest Neighbor Matchings. An adaption to tree like particle arrangements and the inclusion of additional constraints, e.g., incorporating regions of interest are future research objectives.

## 4 Methods

### 4.1 Point Detection

In order to locate the positions of the particles in STED images, we perform point detection via the Python package scikit-image (Walt et al, 2014) (version 0.19.1). This is provided as an optional analysis step in our MultiMatch implementation for the evaluation of intensity matrices.

### 4.2 Interactive Napari Viewer

Multi-color microscopy images, point detection results and MultiMatch output can be loaded into the interactive napari viewer. MultiMatch is compatible with Python package napari (napari contributers, 2019) (version 0.4.18) and an exemplary use-case is described on our repository https://github.com/gnies/multi_match.

### 4.3 Network Flow Implementation

We utilize the minimum-cost flow solver provided in the package ortools (version 9.4.1874) (Perron and Furnon, 2019). For an image containing around 1,000 points in each color channel, a solution of the min cost flow problem can be computed for about 10 different values of *t* in around 1 seconds on a standard laptop. Details on the network architecture and its numerical complexity are given in Supplementary Material A.2.

### 4.4 Simulation Study Setup

In the simulation study discussed in Section 2.2 a predefined number of triplets, pairs, and singlets are generated as follows:

**Step 1:** Draw the coordinate for channel B as *b ∼ U* ([0, 400 *· r*]^2^), where *U* is the continuous uniform distribution.
**Step 2a:** Draw angle *α ∼ U* [0, 2*π*] and normally distributed distance *d_A_∼ N* (*t,* 0.5). Set *a* = *b* (cos(*α*)*d_A_*+ sin(*α*)*d_A_*).
**Step 2b:** Draw *E ∼ N* (0, 0.2) and set angle *β* = *α* + *π* + *E*. Draw *d_C_ ∼ N* (*t,* 0.5) and set *c* = *b* (cos(*β*)*d_C_* + sin(*β*)*d_C_*).
**Step 3:** Round *a, b* and *c* to match the pixel grid [0, 400]^2^ ⊆ 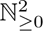

This design favors to simulate triplets of an approximately linear structure. Pairs are simulated by skipping either Step 2a or 2b. Singlets are drawn as in Step 1.

For Section 2.5, quadruples, triplets, pairs, and singlets ***n*** are generated similarly, but replacing and adding

**Step 2b:** Draw angle *β ∼ U* [0, 2*π*] and *d_C_∼ N* (*t,* 0.5) and set *d* = *b* (cos(*β*)*d_C_* + sin(*β*)*d_C_*).
**Step 2c:** Draw angle *γ ∼ U* [0, 2*π*] and *d_D_∼ N* (*t,* 0.5) and set *d* = *c* (cos(*γ*)*d_D_* + sin(*γ*)*d_D_*).

This simulation setup allows arbitrarily curved chain-structures. The distance threshold is always fixed to *t* = 70 nm.

To obtain intensity images close to an experimental STED setup from the simulated point sets we followed the simulation setup introduced in Tameling et al (2021), to mimic experimental STED images of 400 *×* 400 pixels with full-width at half-maximum (FWHM) value of 40 nm (approximately the resolution of the STED microscope) and pixel size 25 nm = 1pixel). In the second simulation study in Section 2.5 the Poisson noise level was on average increased by a factor of 10.

### 4.5 Methods Included in the Simulation Study

For the Ripley’s K based Statistical Object Distance Analysis (SODA, Lagache et al, 2018) we used the triplet colocalization protocol SODA 3 Colors in ICY (version 2.4.0.0, de Chaumont et al, 2012). For the analysis we used default input parameters and set scale threshold per channel to be 100. The plugin BlobProb (Fletcher et al, 2010) was called in ImageJ/Fiji (version 2.3.0/1.53q, Schindelin et al, 2012) and the number of colocalized blobs were considered. We set voxel size to 25 nm in every dimension and the threshold per channel to 100. The ConditionalColoc (Vega-Lugo et al, 2022) from GitHub (https://github.com/kjaqaman/ConditionalColoc) was executed on MATLAB (version R2023a). Particles were detected using the “point-source detection” algorithm provided via the integrated u-track package (https://github.com/DanuserLab/u-track).

For all implementations but ConditionalColoc the detected chain-structure abundances were output as integers. Therefore, we scaled abundances, i.e., divided them by the total number of particles detected in channel B. ConditionColoc already aims to output probabilities that are scaled by detected particles per channel, hence no further transformation of the output was performed by us. Since for all simulated Scenarios the same number of particles was generated in every channel, we ensured that both scaling procedures are comparable. The maximal colocalization threshold is set to *t* = 5 pixels = 125 nm throughout all considered methods.

### 4.6 Nanoruler Samples

Custom-made DNA nanoruler samples featuring one, two, or three fluorophore spots, each consisting of 20 fluorophores (Alexa Fluor488, Alexa Fluor594, Star Red), with a distance between the spots of 70 nm, were purchased from Gattaquant - DNA Nanotechnologies (Gräfelfing, Germany). The biotinylated nanorulers were immobilized on a BSA-biotin-neutravidin surface according to the manufacturer’s specifications.

### 4.7 Stimulated Emission Depletion (STED) Super-Resolution Light Microscopy

Image acquisition was done using a quad scanning STED microscope (Abberior Instruments, Göttingen, Germany) equipped with a UPlanSApo 100x/1,40 Oil objective (Olympus, Tokyo, Japan). Excitation of Alexa Fluor 488, Alexa Fluor 594 and Star Red was achieved by laser beams featuring wave lengths of 485 nm, 561 nm and 640 nm nm respectively. For STED imaging, a laser beam with an emission wavelength of 775 nm was applied. For all images, a pixel size of 25 nm was utilized. Except for contrast stretching and increasement of image brightness, no further image processing was applied.

### 4.8 Data and Code Availability

The Python package MultiMatch is available on GitHub repository https://github.com/gnies/multi match. Code and data to create the main and supplementary figures can be accessed via Zenodo archive https://doi.org/10.5281/zenodo.7221879. Scripts were implemented in R (version 4.1.0) and Python (version 3.8.5).

## Supplementary Material

Theoretical framework of the multi-marginal optimal unbalanced transport matching with chain costs and formulation as network flow (Supplementary Material A), statistical inference on labeling efficiencies (Supplementary Material B), comments on the output from our usage of ConditionalColoc (Supplementary Material C), output of MultiMatch Mode II on the experimental STED images (Supplementary Material D), and additional analyis and simulations scenarios for three-color images and four-color images (Supplementary Material E).

## Acknowledgments

We would like to thank Leo Lehmann for his help in software implementation, Jan-Niklas Dohrke for the graphic illustration of the nanorulers in Figure 1C, and Christiane Elgert and Arndt von Haeseler for constructive criticism on the manuscript. J.N. is supported by the Austrian Science Fund (FWF) project number F78 to Arndt von Haeseler. This work was supported by the European Research Council (ERC AdG no. 835102) (to S.J.). G.N., H.L., S.J. and A.M. are supported by the Deutsche Forschungsgemeinschaft (DFG, German Research Foundation) under Germany’s Excellence Strategy–EXC 2067/1-390729940 and H.L., B.S., S.J. and A.M. by the DFG CRC 1456 “Mathematics of Experiment” (Project Number B04, C06).

