## Supplementary Material for "MultiMatch: Geometry-Informed Colocalization in Multi-Color Super-Resolution Microscopy"

MultiMatch: Optimal Matching Colocalization in Multi-Color Super-Resolution Microscopy

### A Optimal Chain-Matching

#### A.1 Unbalanced Multi-Marginal Formulation

In the following we will denote the sets of two-dimensional particle coordinates in the image domain for each of the $k$ color channels as

$$X^{(1)} := \left\{\boldsymbol{x}_l^{(1)}\right\}_{l=1}^{n_1}, \dots, X^{(k)} := \left\{\boldsymbol{x}_l^{(k)}\right\}_{l=1}^{n_k} \subseteq \mathbb{R}^2, \tag{A1}$$

where number of particles $n_j \in \mathbb{N}_{\geq 0}$ for $j \in \{1, \dots, k\}$. For simplicity and related to the considered data in this article, we will only consider the cases $k = 2, 3$ in the following. Generalization to larger $k$ is straight-forward. In a chain-like particle arrangement of the form $\left(\boldsymbol{x}^{(1)}, \dots, \boldsymbol{x}^{(k)}\right)$ with $\boldsymbol{x}^{(j)} \in X^{(j)}$, all neighbors $\boldsymbol{x}^{(j)}, \boldsymbol{x}^{(j+1)}$ have to be closer than the colocalization threshold $t$ and we will denote according tuples as $\mathbf{d}_t^k$-chains:

**Definition 1** ($\mathbf{d}_t^k$-chain) Fix $k \geq 2$. For sets $X^{(1)}, \dots, X^{(k)}$, a distance $\mathbf{d} : \mathbb{R}^2 \times \mathbb{R}^2 \to \mathbb{R}_{\geq 0}$ and a predefined maximal threshold $t \geq 0$ a tuple of $k$ points

$$\left(\boldsymbol{x}^{(1)}, \dots, \boldsymbol{x}^{(k)}\right) \in \mathbb{R}^{2\times k} \quad \text{with} \quad \boldsymbol{x}^{(j)} \in X^{(j)} \text{ for } j \in \{1, \dots, k\} \tag{A2}$$

is a $\mathbf{d}_t^k$-chain, if pairwise point distances along the fixed tuple point order are smaller or equal than $t$, i.e.,

$$\mathbf{d}\left(\boldsymbol{x}^{(j)}, \boldsymbol{x}^{(j+1)}\right) \leq t \quad \text{for} \quad j \in \{1, \dots, k-1\}. \tag{A3}$$

In the context of our colocalization problem, $\mathbf{d}$ is the Euclidean distance (this can easily be generalized), a $\mathbf{d}_t^3$-chain is a *triplet* and a $\mathbf{d}_t^2$-chain a *pair*. For given $t$, we now aim to detect as many $\mathbf{d}_t^k$-chains as possible:

**Definition 2** (Optimal $\mathbf{d}_t^k$-matching) A collection of pairwise disjoint $\mathbf{d}_t^k$-chains is called $\mathbf{d}_t^k$-matching. It is called *optimal* if its number of chains is maximal among all matchings.

Such an optimal $\mathbf{d}_t^k$-matching can be found by utilizing a multi-marginal and unbalanced formulation of OT. For example, if $k = 3$, for each channel $i = 1, 2, 3$, we interpret coordinates of detected particles as support points with mass 1 of a respective discrete distribution. Due to this discrete structure, the resulting optimization problem will be finite-dimensional. Since in our measurements the number of detected particles per channel might differ, we require an unbalanced formulation to compare distributions with different total masses. A wide variety of penalty terms for mass discrepancies has been studied in the literature, see for instance Liero et al (2016). Our problem formulation is closely related to an $\ell^1$-penalty for unmatched particles, see also Le et al (2022). We first consider the problem of finding optimal $d_t^2$-matchings between two point clouds, i.e. $k = 2$. This can be solved via the following optimization problem:

**Definition 3** (Optimal $\mathbf{d}_t^2$-matchings via unbalanced optimal transport) Let $\lambda \in \mathbb{R}_{\geq 0}$, set the cost function

$$c : \mathbb{R}^2 \times \mathbb{R}^2 \to \mathbb{R}_{\geq 0} \cup \{\infty\}, \quad (x_1, x_2) \mapsto \begin{cases} \mathbf{d}(x_1, x_2) - \lambda & \text{if } \mathbf{d}(x_1, x_2) \leq t, \\ +\infty & \text{otherwise,} \end{cases} \tag{A4}$$

and $\boldsymbol{c} \in \mathbb{R}^{n_1 \times n_2}$ the pairwise cost between all points in $X^{(1)}$ and $X^{(2)}$, defined by $c_{i_1,i_2} = c(\boldsymbol{x}_{i_1}^{(1)}, \boldsymbol{x}_{i_2}^{(2)})$. The optimal unbalanced transport problem of interest can now be stated as the following linear program

$$\begin{aligned} \underset{\boldsymbol{\pi} \in \mathbb{R}^{n_1 \times n_2 \times n_3}}{\arg\min} & \sum_{i_1=1}^{n_1} \sum_{i_2=1}^{n_2} c_{i_1 i_2} \pi_{i_1 i_2} \\ \text{s. t.} & \sum_{i_2=1}^{n_2} \pi_{i_1 i_2} \leq 1 \quad \text{for all } i_1 = 1, \dots, n_1 \\ & \sum_{i_1=1}^{n_1} \pi_{i_1 i_2} \leq 1 \quad \text{for all } i_2 = 1, \dots, n_2 \\ & \pi_{i_1 i_2} \geq 0 \quad \text{for all } (i_1, i_2) \in \{1, \dots, n_1\} \times \{1, \dots, n_2\}. \end{aligned} \tag{A5}$$

Entries of an optimal $\boldsymbol{\pi}$ indicate which particles have been matched. The constraints enforces that each particle can at most be part of one matching, but it may also be discarded. By the definition of the cost vector $\boldsymbol{c}$, the solution of Equation (A5) does not match points $\boldsymbol{x}^{(1)}$ and $\boldsymbol{x}^{(2)}$ as soon as they are farther apart than $t$, but for each matching below distance $t$ there is an incentive by the parameter $\lambda$. For $\lambda$ sufficiently large in comparison to $t$ one can show that the solution yields an optimal $\mathbf{d}_t^2$-matching. Among all optimal matchings the above problem prefers one with the lowest sum of pairwise particle distances among matched particles.

We now generalize this to $k = 3$ via a multi-marginal transport problem.

**Definition 4** (Optimal $\mathbf{d}_t^3$-matchings via unbalanced multi-marginal optimal transport) Let $\lambda \in \mathbb{R}_{\geq 0}$, set the cost function

$$c : \mathbb{R}^2 \times \mathbb{R}^2 \times \mathbb{R}^2 \to \mathbb{R}_{\geq 0} \cup \{\infty\},$$
$$(x_1, x_2, x_3) \mapsto \begin{cases} \mathbf{d}(x_1, x_2) + \mathbf{d}(x_2, x_3) - \lambda & \text{if } \mathbf{d}(x_1, x_2) \leq t \wedge \mathbf{d}(x_2, x_3) \leq t, \\ +\infty & \text{otherwise,} \end{cases} \tag{A6}$$

and let $\boldsymbol{c} \in \mathbb{R}^{n_1 \times n_2 \times n_3}$, be the cost tensor between all triplets in $(X^{(1)}, X^{(2)}, X^{(3)})$, defined by $c_{i_1 i_2 i_3} = c(\boldsymbol{x}_{i_1}^{(1)}, \boldsymbol{x}_{i_2}^{(2)}, \boldsymbol{x}_{i_3}^{(3)})$. Then the unbalanced multi-marginal OT problem can be stated as the following linear program:

$$\begin{aligned} \underset{\boldsymbol{\pi} \in \mathbb{R}^{n_1 \times n_2 \times n_3}}{\arg\min} & \sum_{i_1=1}^{n_1} \sum_{i_2=1}^{n_2} \sum_{i_3=1}^{n_3} c_{i_1 i_2 i_3} \pi_{i_1 i_2 i_3} \\ \text{s. t. } & \sum_{i_2=1}^{n_2} \sum_{i_3=1}^{n_3} \pi_{i_1 i_2 i_3} \leq 1 \quad \text{for all } i_1 \in [n_1] \\ & \sum_{i_1=1}^{n_1} \sum_{i_3=1}^{n_3} \pi_{i_1 i_2 i_3} \leq 1 \quad \text{for all } i_2 \in [n_2] \\ & \sum_{i_1=1}^{n_1} \sum_{i_3=1}^{n_3} \pi_{i_1 i_2 i_3} \leq 1 \quad \text{for all } i_3 \in [n_1] \\ & \pi_{i_1 i_2 i_3} \geq 0 \quad \text{for all } (i_1, i_2, i_3) \in [n_1] \times [n_2] \times [n_3], \end{aligned} \tag{A7}$$

where we used the notation $[n] = \{1, \ldots, n\}$. As above note that per the marginal constraints particles may be matched at most once and can also be discarded. Likewise, by definition of the cost vector $\boldsymbol{c}$ only allows matchings between points that are valid $\mathbf{d}_t^3$-chains. Analogously there is a matching incentive via the parameter $\lambda$ and for sufficiently high values (relative to $t$) one can show that the above problem provides an optimal $\mathbf{d}_t^3$-matching. Among all these matchings, one with minimal sum of pairwise distances is selected by the problem.

Generalization of Definition 4 to arbitrary $k$ is now obvious, leading to a multi-marginal problem with $k$ marginals. In general, multi-marginal problems quickly become numerically impractical due to the large number of variables. The cost function $c$ in (A6) has a chain structure, i.e. it can be written as a sum of functions only depending on $(x_1, x_2)$ and $(x_2, x_3)$. This chain structure allows the reformulation of the problem as a much more compact network flow problem (see Section below), and it implies the existence of optimal binary matchings. Problems where the cost exhibits a tree-structure can still be solved efficiently, see Beier et al (2022) and references therein, but they cannot be formulated as network flow problems and do not exhibit binary minimizers in general.

#### A.2 Network Flow Formulation of MultiMatch

In this section we show that the multi-marginal optimal unbalanced transport problem corresponds to a min cost flow problem if the cost function has a chain structure as in (A6). This has two relevant consequences:

1. It guarantees that (A7) has integer solutions and thus indeed corresponds to a matching problem, which in general does not hold true for discrete OT problems;
2. It allows us to solve the multi-marginal optimal unbalanced transport problem efficiently.

**Definition 5** Let $(V, E)$ be a directed graph with a source node $S \in V$, a target node $T \in V$, an edge capacity function $l_E : E \to \mathbb{R} \cup \infty$ and an edge cost function $c_E : E \to \mathbb{R} \cup \infty$. Then we call $(V, E, c_E, l_E)$ a flow network. Given an amount of flow, $m \in \mathbb{R}_+$ the min cost flow problem consists in finding a function $f : E \to \mathbb{R}$ that solves the following optimization problem:

$$
\begin{aligned}
&\min_f \sum_{(u,v)\in E} f(u,v) c_E(u,v) \\
&\text{s.t.} \quad 0 \le f(u,v) \le l_E(u,v) \text{ for all } (u,v) \in E && \text{(capacity constraints)} \\
&\sum_{\{u:(u,v)\in E\}} f(u,v) - \sum_{\{w:(v,w)\in E\}} f(v,w) = 0 \text{ for all } v \neq S, T && \text{(flow conservation)} \\
&\sum_{\{u:(S,u)\in E\}} f(S,u) - \sum_{\{v:(v,S)\in E\}} f(v,S) = m && \text{(flow source)} \\
&\sum_{\{u:(u,T)\in E\}} f(u,T) - \sum_{\{v:(T,v)\in E\}} f(T,v) = m && \text{(flow sink).}
\end{aligned}
$$

Notably, due to the total unimodularity of the constraint matrix, the min cost flow problem with integer total flow $m$ and integer capacity function $l_E$ has an integer solution (Theorem 13.11 in Alexander (2003)). In the following, we recast (A7) to a min cost flow problem (see sketch in Figure A1):

- **Node set** $V$**:** Define source node $S \in V$ and target node $T \in V$ and add two nodes $v_l^{(j)}$ and $\hat{v}_l^{(j)}$ for each detected particle position $\mathbf{x_l^{(j)}}$ in Equation (A1).
- **Edge set** $E$**:**
  - Connect nodes referring to the same detected point and set edge costs $c_E(v_l^{(j)}, \hat{v}_l^{(j)}) = -\frac{\lambda}{k}$ where $k$ is the number of point clouds as in (A1).
  - Add all possible edges of form $(\hat{v}^{(j)}, v^{(j+1)}) \in E$ for $j = 1, \ldots, k-1$. Set edge costs

$$
c_E(\hat{v}^{(j)}, v^{(j+1)}) = \begin{cases} \infty, & \text{if } \boldsymbol{d}(\boldsymbol{x}^{(j)}, \boldsymbol{x}^{(j+1)}) > t \\ \boldsymbol{d}(\boldsymbol{x}^{(j)}, \boldsymbol{x}^{(j+1)}), & \text{otherwise.} \end{cases}
$$

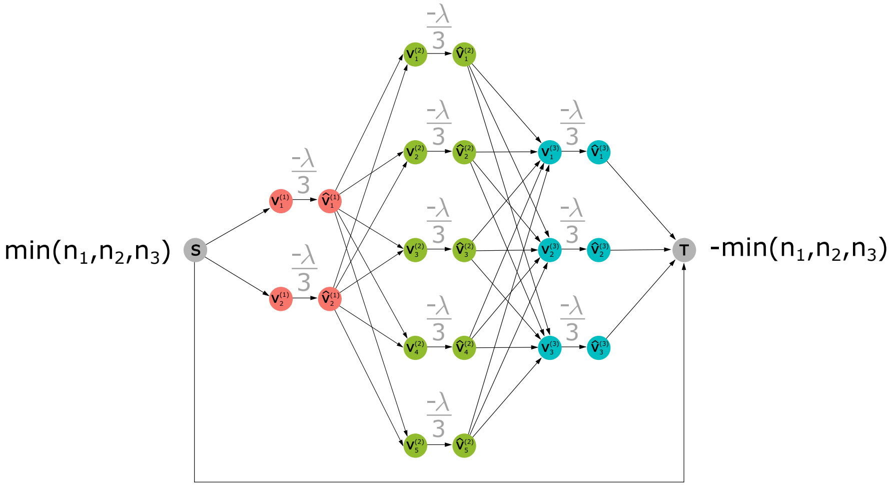


**Fig. A1** Scheme of the implemented min cost network flow problem as explained in Section A.2.

– Include source and target nodes via edges of form $(S, v^{(1)}), (\hat{v}^{(k)}, T), (S, T) \in E$, and set its costs to 0.
– Define edge capacities

$$l_E(v_i, v_j) = \begin{cases} \infty, & \text{if } v_i = S \text{ and } v_j = T \\ 1, & \text{otherwise.} \end{cases}$$

**Proposition 1** *Let $f : E \to \mathbb{R}$ be an integer solution of the min cost flow problem for the flow network $(V, E, c_E, l_E)$ defined above with transported mass $m = \min(n_1, n_2, n_3)$. Then one of the optimal solutions $\boldsymbol{\pi}^*$ of the multi-marginal optimal unbalanced transport problem (A7) is given by,*

$$\pi^*_{i_1 i_2 i_3} = f(\hat{v}^{(1)}_{i_1}, v^{(2)}_{i_2}) f(\hat{v}^{(2)}_{i_2}, v^{(3)}_{i_3}), \tag{A8}$$

*for $i_1 \in [n_1], i_2 \in [n_2]$ and $i_3 \in [n_3]$ with notation $[n] = \{1, \dots, n\}$.*

*Proof* First we show that $\boldsymbol{\pi}^*$ as defined in (A8) is in fact a valid transport plan for (A7). For any $i_3 \in [n_3]$ we have that, using the conservation constraint,

$$\begin{aligned} \sum_{i_1=1}^{n_1} \sum_{i_2=1}^{n_2} \pi^*_{i_1 i_2 i_3} &= \sum_{i_2=1}^{n_2} f(\hat{v}^{(2)}_{i_2}, v^{(3)}_{i_3}) \left( \sum_{i_1=1}^{n_2} f(\hat{v}^{(1)}_{i_1}, v^{(2)}_{i_2}) \right) \\ &= \sum_{i_2=1}^{n_2} f(\hat{v}^{(2)}_{i_2}, v^{(3)}_{i_3}) f(\hat{v}^{(2)}_{i_2}, v^{(2)}_{i_2}) \\ &\leq \sum_{i_2=1}^{n_2} f(\hat{v}^{(2)}_{i_2}, v^{(3)}_{i_3}) = f(\hat{v}^{(3)}_{i_3}, v^{(3)}_{i_3}) \leq 1 \end{aligned}$$

Analogously it is easy to verify that $\boldsymbol{\pi}^*$ satisfies

$$\sum_{i_1=1}^{n_1} \sum_{i_3=1}^{n_3} \pi^*_{i_1 i_2 i_3} \leq 1 \quad \text{for all } i_2 \in [n_2]$$
$$\sum_{i_2=1}^{n_2} \sum_{i_3=1}^{n_3} \pi^*_{i_1 i_2 i_3} \leq 1 \quad \text{for all } i_1 \in [n_1].$$

Hence, $\boldsymbol{\pi}^*$ is a feasible solution of (A7). Further, since the source node $S$ is directly connected to the target node $T$ with an edge of infinite capacity and finite cost, the total flow cost must be finite. This implies that for any $i_1 \in [n_1], i_2 \in [n_2]$ and $i_3 \in [n_3]$, we have that $f(\hat{v}^{(1)}_{i_1}, v^{(2)}_{i_2}) = 0$ if $\boldsymbol{d}(\boldsymbol{x}^{(1)}_{i_1}, \boldsymbol{x}^{(2)}_{i_2}) > t$ and $f(\hat{v}^{(2)}_{i_2}, v^{(3)}_{i_3}) = 0$ if $\boldsymbol{d}(\boldsymbol{x}^{(2)}_{i_2}, \boldsymbol{x}^{(3)}_{i_3}) > t$. Hence, using the shorthand notation

$$\langle \boldsymbol{c}, \boldsymbol{\pi} \rangle = \sum_{i_1=1}^{n_1} \sum_{i_2=1}^{n_2} \sum_{i_3=1}^{n_3} c_{i_1 i_2 i_3} \pi_{i_1 i_2 i_3},$$

we can rewrite the total cost of the transport problem as

$$\langle \boldsymbol{c}, \boldsymbol{\pi}^* \rangle = \sum_{i_1=1}^{n_1} \sum_{i_2=1}^{n_2} \sum_{i_3=1}^{n_3} \Big( \boldsymbol{d}(\boldsymbol{x}^{(1)}_{i_1}, \boldsymbol{x}^{(2)}_{i_2}) + \boldsymbol{d}(\boldsymbol{x}^{(2)}_{i_2}, \boldsymbol{x}^{(3)}_{i_3}) - \lambda \Big) f(\hat{v}^{(1)}_{i_1}, v^{(2)}_{i_2}) f(\hat{v}^{(2)}_{i_2}, v^{(3)}_{i_3}).$$

By the flow conservation constraints and the fact that $f$ is an integer solution, we can simply reformulate the sum above in terms of the network flow cost function to obtain

$$\langle \boldsymbol{c}, \boldsymbol{\pi}^* \rangle = \sum_{(u,v) \in E} c_E(u,v) f(u,v).$$

Let us now assume that there exists a feasible solution of (A7), $\tilde{\boldsymbol{\pi}}$, such that

$$\langle \boldsymbol{c}, \tilde{\boldsymbol{\pi}} \rangle < \langle \boldsymbol{c}, \boldsymbol{\pi}^* \rangle.$$

Then we can define the flow $\tilde{f} : E \to \mathbb{R}$ by setting:

$$\tilde{f}(\hat{v}^{(1)}_{i_1}, v^{(2)}_{i_2}) = \sum_{i_3=1}^{n_3} \tilde{\pi}_{i_1 i_2 i_3} \text{ and } \tilde{f}(\hat{v}^{(2)}_{i_2}, v^{(3)}_{i_3}) = \sum_{i_1=1}^{n_1} \tilde{\pi}_{i_1 i_2 i_3},$$

for $i_1 \in [n_1], i_2 \in [n_2]$ and $i_3 \in [n_3]$. The value of the flow on the remaining nodes of $E$ can then be determined by the conservation constraints. In particular, we have $\tilde{f}(S,T) = \min\{n_1, n_2, n_3\} - \sum_{i_1=1}^{n_1} \sum_{i_2=1}^{n_2} \sum_{i_3=1}^{n_3} \pi_{i_1 i_2 i_3}$. This flow is a feasible solution of the given min cost flow problem and hence, by the definition of the cost function for the edges we can derive a contradiction:

$$\sum_{(u,v) \in E} c_E(u,v) \tilde{f}(u,v) = \langle \boldsymbol{c}, \tilde{\boldsymbol{\pi}} \rangle < \sum_{(u,v) \in E} c_E(u,v) f(u,v).$$

□

As a result of Proposition 1, we immediately obtain that the multi-marginal optimal unbalanced transport problem (A7) has an integer solution and hence provides one-to-one point matchings.

Another significant consequence of Proposition 1 is that we can solve the unbalanced optimal transport problem given in (A7) efficiently. While it is often unfeasible to compute directly the solution of the $n_1 \cdot n_2 \cdots n_k$-dimensional linear programming problem in (A7), the min cost flow problem can be solved by the Scaling Minimum-Cost Flow Algorithm in Goldberg (1997) in $O(|V|^2|E|\log(|V|))$ elementary operations, where $|V|$ is the number of nodes, $|E|$ is the number of edges. In our case the number of nodes is of the order $O(n_1 \cdot \cdots \cdot n_k)$ and the number of edges can be upper bounded by an expression of the order $O(n_1 \cdot n_2^2 \cdot \cdots \cdot n_{k-1}^2 \cdot n_k)$. In practice, it is further possible to omit all edges with infinite cost, since the source $S$ and the sink $T$ are connected through an edge of cost 0 and with infinite capacity. This implies that for small $t$ much fewer edges to the network are added which results in better computational performance.

### B The probability of Correct Tuple Detection

The quality of fluorescence microscopy suffers from non-optimal labeling efficiencies and point detection errors. This will be adressed by a statistical framework to infer on how many of the detected structures in the image actually concur with the ground truth biological structure and how many detections represent only incomplete parts of the underlying particle assembly. For color channels $i \in \{1, \ldots, k\}$ let

$$\left\{\xi_j^{(i)}\right\}_{j=1}^{n_i} \subset \mathbb{R}^2 \tag{B9}$$

be the pairs of coordinates of all particles that lie within the scope of the microscope. Note that these point clouds do not necessarily equal those defined in (A1) describing the coordinates of detected particles, since we might not be able to measure all of the existing particles to do unsuccessful labeling or point detection errors.

**Definition 6** (Labeling Efficiency) For each color channel $i \in \{1, \ldots, k\}$ we assume that there is a specific probability $s_i \in (0, 1]$ quantifying whether a particle of this channel is successfully imaged and detected. For simplicity in the following we will always call probabilities $s_i$ *labeling efficiencies.*

We further assume that the random event of successful detection is statistically independent for each point. Accordingly, the detection success can be described by independent Bernoulli variables

$$\left\{Z_j^{(i)}\right\}_{j=1}^{n_i} \sim \mathrm{Ber}(s_i), \tag{B10}$$

where $s_i \in (0, 1]$ and $\xi_j^{(i)}$ is detectable, if and only if $Z_j^{(i)} = 1$.

If there exists a true $\mathbf{d}_t^k$-chain of form $\left(\xi^{(1)}, \ldots, \xi^{(k)}\right)$, then this can only be correctly identified as such, if each of the included particles was detected, i.e., if and only if $\prod_{i=1}^{k} Z^{(i)} = 1$. From independence it follows that

$$\prod_{i=1}^{k} Z^{(i)} \sim \mathrm{Ber}\left(\prod_{i=1}^{k} s_i\right). \tag{B11}$$

#### B.1 Estimating the true abundances of structures

Detecting an ABC triplet correctly is $\mathrm{Ber}(s_A s_B s_C)$ distributed. Therefore, all possible substructures that can be detected conditioned on the true underlying ABC triplet, i.e.,

1. ABC triplet, if we see all particles
2. AB pair, if we do not see C
3. BC pair, if we do not see A
4. *AC substructure, if we do not see B – which is detected as A and C singlets*
5. A singlet, if we do not see B and C
6. B singlet, if we do not see A and C
7. C singlet, if we do not see A and B
8. $\emptyset$, *if we do not see A,B and C– which can not be detected at all,*

can accordingly be modeled as Multinomial random variable

$$\boldsymbol{W}_{\cdot|ABC} = \begin{bmatrix} W_{ABC|ABC} \\ W_{AB|ABC} \\ W_{BC|ABC} \\ W_{AC|ABC} \\ W_{A|ABC} \\ W_{B|ABC} \\ W_{C|ABC} \\ W_{\emptyset|ABC} \end{bmatrix}. \tag{B12}$$

This can be done in the same manner for all other structures of interest, i.e., true AB and BC pairs and A,B and C singlets (and their respective substructures) yielding random variables $W_{\cdot|AB}, W_{\cdot|BC}, W_{\cdot|A}, W_{\cdot|B}, W_{\cdot|C}$. The actual detectable numbers of those structures are

$$\begin{aligned} W_{ABC} &= \sum W_{ABC|\cdot}, \\ W_{AB} &= \sum W_{AB|\cdot}, \\ W_{BC} &= \sum W_{BC|\cdot}, \\ W_{A} &= \sum W_{A|\cdot} + \sum W_{AC|\cdot}, \\ W_{B} &= \sum W_{B|\cdot}, \\ W_{C} &= \sum W_{C|\cdot} + \sum W_{AC|\cdot}, \end{aligned} \tag{B13}$$

which define a random variable $\boldsymbol{W} = (W_{ABC}, W_{AB}, W_{BC}, W_A, W_B, W_C)^T$. This leads to a statistical framework, that allows us to estimate the true underlying structures abundances from the detected number of structures.

**Theorem 2** *Let known, positive labeling efficiencies $s_A > 0, s_B > 0$ and $s_C > 0$ and unknown structure abundances $\boldsymbol{n} = (n_{ABC}, n_{AB}, n_{BC}, n_A, n_B, n_C)^T$ and define $N = \sum_{i \in \{ABC,\dots,C\}} n_i$. Assume the multinomial model as described in Equation (B12) and Equation (B13).*

***Part 1:*** *An unbiased estimator $\hat{\boldsymbol{n}}$ of true abundances $\boldsymbol{n}$ is given as*

$$\hat{\boldsymbol{n}} = \begin{bmatrix} \frac{1}{s_A s_B s_C} & 0 & 0 & 0 & 0 & 0 \\ \frac{s_C - 1}{s_A s_B s_C} & \frac{1}{s_A s_B} & 0 & 0 & 0 & 0 \\ \frac{s_A - 1}{s_A s_B s_C} & 0 & \frac{1}{s_B s_C} & 0 & 0 & 0 \\ \frac{s_B - 1}{s_A s_B} & \frac{s_B - 1}{s_A s_B} & 0 & \frac{1}{s_A} & 0 & 0 \\ \frac{(1-s_A)(1-s_C)}{s_A s_B s_C} & \frac{s_A - 1}{s_A s_B} & \frac{s_C - 1}{s_B s_C} & 0 & \frac{1}{s_B} & 0 \\ \frac{s_B - 1}{s_B s_C} & 0 & \frac{s_B - 1}{s_B s_C} & 0 & 0 & \frac{1}{s_C} \end{bmatrix} \boldsymbol{W}. \tag{B14}$$

***Part 2:*** *For $\boldsymbol{n} \to \infty$ entrywise, $n_j/N \to f_j$ with $\infty > f_j > 0$ constant for each $j \in \{ABC, \dots, C\}$, and $\Theta\Sigma(\hat{\boldsymbol{n}})\Theta^T$ invertible,*

$$P\left(\Xi \leq \chi^2_{6,\alpha}\right) \leq 1 - \alpha, \tag{B15}$$

*where*

$$\Xi = (\hat{\boldsymbol{n}} - \boldsymbol{n})^T (\Theta\mu)^T \left(\Theta\Sigma(\hat{\boldsymbol{n}})\Theta^T\right)^{-1} (\Theta\mu)(\hat{\boldsymbol{n}} - \boldsymbol{n}) \tag{B16}$$

*and $\chi^2_{6,\alpha}$ is the $\alpha$-quantile of a chi-squared distribution with 6 degrees of freedom and with $\Theta$, $\mu$ and $\Sigma(\hat{\boldsymbol{n}})$ defined as in the following proof. If $\left(\Theta\Sigma(\hat{\boldsymbol{n}})\Theta^T\right)^{-1}$ does not exist, we get Equation (B15) with $\chi^2_{r,\alpha}$ plugging its pseudoinverse $\left(\Theta\Sigma(\hat{\boldsymbol{n}})\Theta^T\right)^{+}$ in Equation (B16), where $r = \operatorname{rank}\left(\Theta\Sigma(\hat{\boldsymbol{n}})\Theta^T\right)$.*

*Proof* **Part 1:** Conditioned on a true ABC triplet, the number of (mis)specifications resulting from incomplete labeling efficiencies is multinomially distributed:

$$\boldsymbol{W}_{\cdot|ABC} = \begin{bmatrix} W_{ABC|ABC} \\ W_{AB|ABC} \\ W_{BC|ABC} \\ W_{AC|ABC} \\ W_{A|ABC} \\ W_{B|ABC} \\ W_{C|ABC} \\ W_{\emptyset|ABC} \end{bmatrix} \sim \mathrm{Mnom}(n_{ABC}, \boldsymbol{p}_{ABC}) \tag{B17}$$

with probability vector

$$\boldsymbol{p}_{ABC} = \begin{bmatrix} s_A s_B s_C \\ s_A s_B (1-s_C) \\ (1-s_A) s_B s_C \\ s_A (1-s_B) s_C \\ s_A (1-s_B)(1-s_C) \\ (1-s_A) s_B (1-s_C) \\ (1-s_A)(1-s_B) s_C \\ (1-s_A)(1-s_B)(1-s_C) \end{bmatrix}, \tag{B18}$$

where $\sum_{j=1}^{8} \boldsymbol{p}_{ABC}[j] = 1$. Accordingly, the abundances of (mis)detections of a true AB pair are

$$\begin{bmatrix} W_{ABC|AB} \\ W_{AB|AB} \\ W_{BC|AB} \\ W_{AC|AB} \\ W_{A|AB} \\ W_{B|AB} \\ W_{C|AB} \\ W_{\emptyset|AB} \end{bmatrix} \sim \text{Mnom}(n_{AB}, \boldsymbol{p}_{AB}) \tag{B19}$$

with

$$\boldsymbol{p}_{AB} = \begin{bmatrix} 0 \\ s_A s_B \\ 0 \\ 0 \\ s_A (1-s_B) \\ (1-s_A) s_B \\ 0 \\ (1-s_A)(1-s_B) \end{bmatrix}. \tag{B20}$$

This can be done accordingly for all other stuctures of interest, i.e. BC pairs and A,B and C singlets yielding

$$\begin{aligned} \boldsymbol{W}_{\cdot|ABC} &\sim \text{Mnom}(n_{ABC}, \boldsymbol{p}_{ABC}) \\ \boldsymbol{W}_{\cdot|AB} &\sim \text{Mnom}(n_{AB}, \boldsymbol{p}_{AB}) \\ \boldsymbol{W}_{\cdot|BC} &\sim \text{Mnom}(n_{BC}, \boldsymbol{p}_{BC}) \\ \boldsymbol{W}_{\cdot|A} &\sim \text{Mnom}(n_{A}, \boldsymbol{p}_{A}) \\ \boldsymbol{W}_{\cdot|B} &\sim \text{Mnom}(n_{B}, \boldsymbol{p}_{B}) \\ \boldsymbol{W}_{\cdot|C} &\sim \text{Mnom}(n_{C}, \boldsymbol{p}_{C}) \end{aligned} \tag{B21}$$

with

$$\boldsymbol{p}_{BC} = \begin{bmatrix} 0 \\ 0 \\ s_B s_C \\ 0 \\ 0 \\ s_B (1-s_C) \\ (1-s_B) s_C \\ (1-s_B)(1-s_C) \end{bmatrix}, \boldsymbol{p}_{A} = \begin{bmatrix} 0 \\ 0 \\ 0 \\ 0 \\ s_A \\ 0 \\ 0 \\ (1-s_A) \end{bmatrix}, \boldsymbol{p}_{B} = \begin{bmatrix} 0 \\ 0 \\ 0 \\ 0 \\ 0 \\ s_B \\ 0 \\ (1-s_B) \end{bmatrix}, \boldsymbol{p}_{C} = \begin{bmatrix} 0 \\ 0 \\ 0 \\ 0 \\ 0 \\ 0 \\ s_C \\ (1-s_C) \end{bmatrix}. \tag{B22}$$

Note, that $\emptyset$ can not be detected at all and substructure AC is counted as a separate A and C singlet Hence, the total numbers of detected triplets, pairs and singlets are defined as the following sums

$$\begin{aligned} W_{ABC} &= W_{ABC|ABC} \\ W_{AB} &= W_{AB|ABC} + W_{AB|AB} \\ W_{BC} &= W_{BC|ABC} + W_{BC|BC} \\ W_{A} &= W_{A|ABC} + W_{AC|ABC} + W_{A|AB} + W_{A|A} \\ W_{B} &= W_{B|ABC} + W_{B|AB} + W_{B|BC} + W_{B|B} \\ W_{C} &= W_{C|ABC} + W_{AC|ABC} + W_{C|BC} + W_{C|C}. \end{aligned} \tag{B23}$$

This can be rewritten as

$$\boldsymbol{W} = \begin{bmatrix} W_{ABC} \\ W_{AB} \\ W_{BC} \\ W_{A} \\ W_{B} \\ W_{C} \end{bmatrix} = \Theta \left( \boldsymbol{W}_{\cdot|ABC} + \boldsymbol{W}_{\cdot|AB} + \boldsymbol{W}_{\cdot|BC} + \boldsymbol{W}_{\cdot|A} + \boldsymbol{W}_{\cdot|B} + \boldsymbol{W}_{\cdot|C} \right), \tag{B24}$$

using the transformation matrix

$$\Theta = \begin{bmatrix} 1 & 0 & 0 & 0 & 0 & 0 & 0 & 0 \\ 0 & 1 & 0 & 0 & 0 & 0 & 0 & 0 \\ 0 & 0 & 1 & 0 & 0 & 0 & 0 & 0 \\ 0 & 0 & 0 & 1 & 1 & 0 & 0 & 0 \\ 0 & 0 & 0 & 0 & 0 & 1 & 0 & 0 \\ 0 & 0 & 0 & 1 & 0 & 0 & 1 & 0 \end{bmatrix} \in \mathbb{R}^{6\times 8}. \tag{B25}$$

With this definition of $\Theta$ we delete the last entry in each binomial distributed vector and add an AC substructure appearance to singlet detections A and B. By Equation (B24) we get that

$$\mathbb{E}[\boldsymbol{W}] = \Theta \mu \boldsymbol{n} \tag{B26}$$

with

$$\mu = \left[ \boldsymbol{p}_{ABC} \ \boldsymbol{p}_{AB} \ \boldsymbol{p}_{BC} \ \boldsymbol{p}_{A} \ \boldsymbol{p}_{B} \ \boldsymbol{p}_{C} \right] \in \mathbb{R}^{8\times 6}. \tag{B27}$$

Hence, with positive labelling efficiencies $s_A > 0, s_B > 0$ and $s_C > 0$, multiplying

$$(\Theta\mu)^{-1} = \begin{bmatrix} \frac{1}{s_A s_B s_C} & 0 & 0 & 0 & 0 & 0 \\ \frac{s_C - 1}{s_A s_B s_C} & \frac{1}{s_A s_B} & 0 & 0 & 0 & 0 \\ \frac{s_A - 1}{s_A s_B s_C} & 0 & \frac{1}{s_B s_C} & 0 & 0 & 0 \\ \frac{s_B - 1}{s_A s_B} & \frac{s_B - 1}{s_A s_B} & 0 & \frac{1}{s_A} & 0 & 0 \\ \frac{(1-s_A)(1-s_C)}{s_A s_B s_C} & \frac{s_A - 1}{s_A s_B} & \frac{s_C - 1}{s_B s_C} & 0 & \frac{1}{s_B} & 0 \\ \frac{s_B - 1}{s_B s_C} & 0 & \frac{s_B - 1}{s_B s_C} & 0 & 0 & \frac{1}{s_C} \end{bmatrix} \tag{B28}$$

with $\boldsymbol{W}$ introduces an unbiased estimator $\hat{\boldsymbol{n}}$.

**Part 2:** We utilize that by the central limit theorem for a multinomially distributed random variable $\boldsymbol{M} \sim \text{Mnom}(m, \boldsymbol{p})$ with probability vector $\boldsymbol{p} = (p_1, p_2, \ldots, p_k)^T$

$$\frac{1}{\sqrt{m}} (\boldsymbol{M} - m\boldsymbol{p}) \xrightarrow{\mathcal{D}} \mathcal{N}_k \left( \boldsymbol{0}_k, \text{diag}(\boldsymbol{p}) - \boldsymbol{p}\boldsymbol{p}^T \right) \quad \text{for} \quad m \to \infty, \tag{B29}$$

where

$$\operatorname{diag}(\boldsymbol{p}) = \begin{bmatrix} p_1 & 0 & \cdots \\ 0 & p_2 & \cdots \\ \vdots & \vdots & p_k \end{bmatrix} \tag{B30}$$

and $\mathbf{0}_k = (0, ..., 0)^T \in \mathbb{R}^k$ (see, e.g., Morris, 1975). Hence, for $\boldsymbol{n}$ entrywise large enough, we can approximate properly scaled independent, multinomial random vectors

$$\boldsymbol{W}_{\cdot|ABC}, \boldsymbol{W}_{\cdot|AB}, \boldsymbol{W}_{\cdot|BC}, \boldsymbol{W}_{\cdot|A}, \boldsymbol{W}_{\cdot|B}, \boldsymbol{W}_{\cdot|C} \tag{B31}$$

with multi-dimensional normal distributions, respectively. In the following assume $\boldsymbol{n} \to \infty$ entrywise and $n_j/N \to f_j$ with $\infty > f_j > 0$ constant for each $j \in \{ABC, \ldots, C\}$, where $N = \sum_{i \in \{ABC,\ldots,C\}} n_i$. Then, it holds that

$$\begin{aligned} \sum_{i\in\{ABC,\ldots,C\}} \sqrt{\frac{n_i}{N}} \frac{1}{\sqrt{n_i}} \left(\boldsymbol{W}_{\cdot|i} - n_i \boldsymbol{p}_i\right) &= \sqrt{\frac{1}{N}} \sum_{i\in\{ABC,\ldots,C\}} \left(\boldsymbol{W}_{\cdot|i} - n_i \boldsymbol{p}_i\right) \\ \xrightarrow{\mathcal{D}} \mathcal{N}_8 \left( \mathbf{0}_8, \sum_{i\in\{ABC,\ldots,C\}} f_i \left(\operatorname{diag}(\boldsymbol{p}_i) - \boldsymbol{p}_i \boldsymbol{p}_i^T\right)\right). \end{aligned} \tag{B32}$$

For now, suppose $\sum n_i \left(\operatorname{diag}(\boldsymbol{p}_i) - \boldsymbol{p}_i\boldsymbol{p}_i^T\right)$ is invertible. Then in the limit

$$\begin{aligned} &\left(\sum f_i \left(\operatorname{diag}(\boldsymbol{p}_i) - \boldsymbol{p}_i \boldsymbol{p}_i^T\right)\right)^{-1/2} \sqrt{\frac{1}{N}} \sum \left(\boldsymbol{W}_{\cdot|i} - n_i \boldsymbol{p}_i\right) \\ =& \left(\sum N f_i \left(\operatorname{diag}(\boldsymbol{p}_i) - \boldsymbol{p}_i \boldsymbol{p}_i^T\right)\right)^{-1/2} \sum \left(\boldsymbol{W}_{\cdot|i} - n_i \boldsymbol{p}_i\right) \\ =& \left(\sum n_i \left(\operatorname{diag}(\boldsymbol{p}_i) - \boldsymbol{p}_i \boldsymbol{p}_i^T\right)\right)^{-1/2} \sum \left(\boldsymbol{W}_{\cdot|i} - n_i \boldsymbol{p}_i\right) \end{aligned} \tag{B33}$$

and hence

$$\left(\sum n_i \left(\operatorname{diag}(\boldsymbol{p}_i) - \boldsymbol{p}_i \boldsymbol{p}_i^T\right)\right)^{-1/2} \sum \left(\boldsymbol{W}_{\cdot|i} - n_i \boldsymbol{p}_i\right) \xrightarrow{\mathcal{D}} \mathcal{N}_8\left(\mathbf{0}_8, I_{8\times 8}\right), \tag{B34}$$

where $I_{8\times8}$ is the 8-dimensional identity matrix. In the following we denote

$$\Sigma(\boldsymbol{n}) = \left(\sum n_i \left(\operatorname{diag}(\boldsymbol{p}_i) - \boldsymbol{p}_i \boldsymbol{p}_i^T\right)\right). \tag{B35}$$

Multiplying $(\Theta\mu)^{-1}\Theta$ with Equation (B32) consequently yields

$$\begin{aligned} &\left((\Theta\mu)^{-1}\Theta\Sigma(\boldsymbol{n})\Theta^T \left((\Theta\mu)^{-1}\right)^T\right)^{-1/2} \left((\Theta\mu)^{-1}\Theta \sum \boldsymbol{W}_{\cdot|i} - (\Theta\mu)^{-1}\Theta \sum n_i \boldsymbol{p}_i\right) \\ =& \left((\Theta\mu)^{-1}\Theta\Sigma(\boldsymbol{n})\Theta^T \left((\Theta\mu)^{-1}\right)^T\right)^{-1/2} (\hat{\boldsymbol{n}} - \boldsymbol{n}) \xrightarrow{\mathcal{D}} \mathcal{N}_6\left(\mathbf{0}_6, I_{6\times 6}\right) \end{aligned} \tag{B36}$$

with $\hat{\boldsymbol{n}} = (\Theta\mu)^{-1}\Theta \sum \boldsymbol{W}_{\cdot|i}$ and $\boldsymbol{n} = (\Theta\mu)^{-1}\Theta\mu\boldsymbol{n} = (\Theta\mu)^{-1}\Theta \sum n_i \boldsymbol{p}_i$. By law of large numbers, it holds that

$$\frac{1}{N}(\hat{\boldsymbol{n}} - \boldsymbol{n}) = \frac{\hat{\boldsymbol{n}}}{N} - \frac{\boldsymbol{n}}{N} \xrightarrow{\mathcal{P}} \mathbf{0}_6. \tag{B37}$$

and hence for all $j \in \{ABC, \ldots, C\}$

$$\frac{\hat{n}_j}{N} \xrightarrow{\mathcal{P}} f_j. \tag{B38}$$

By Slutsky's Lemma we can use Equation (B38) to replace $\boldsymbol{n}$ in $\Sigma(\boldsymbol{n})$ with $\hat{\boldsymbol{n}}$. For $\boldsymbol{n} \to \infty$ entrywise this yields

$$\Xi = (\hat{\boldsymbol{n}} - \boldsymbol{n})^T (\Theta\mu)^T \left(\Theta\Sigma(\hat{\boldsymbol{n}})\Theta^T\right)^{-1} (\Theta\mu)(\hat{\boldsymbol{n}} - \boldsymbol{n}) \xrightarrow{\mathcal{D}} \chi^2_6. \tag{B39}$$

In case $\Theta\Sigma(\hat{\boldsymbol{n}})\Theta^T$ is not invertable, one can use its pseudoinverse yielding convergence to a chi-square distribution with $r$ degrees of freedom, i.e., $\chi^2_r$ in Equation (B39), where $r = \text{rank}\left(\Theta\Sigma(\hat{\boldsymbol{n}})\Theta^T\right)$. □

With Part 2 of Theorem 2 we can construct a confidence ellipsoid around $\hat{\boldsymbol{n}}$ in a straight-forward manner. To show that $\Xi$ in our setting is approximately chi-square distributed for finite sample sizes and to compare simulated and theoretical coverages of $\hat{\boldsymbol{n}}$, we performed a simulation study as described in the following section.

#### B.2 Simulation study of incomplete labeling efficiencies

We simulated incomplete labeling efficiencies by following the statistical framework developed in the Proof of Theorem 2, Part 1: The numbers of detectable triplets, pairs and singlets $\boldsymbol{W}$ were simulated from true abundances $\boldsymbol{n}$ by drawing 10,000 values from respective multinomial distributions based on predefined staining efficiencies $s_A, s_B, s_C$ (see multinomial model in Equation (B12) and Equation (B13)). All combinations of abundances and staining efficiencies that were simulated are listed in Table B1, where we also recorded the respective empirical coverage of constructed joint confidence ellipsoids at a theoretical coverage of $1 - \alpha = 0.90$.

For $s_A = s_B = s_C = 0.95$ and $n_{ABC} = 500, n_A = n_B = s_C = 50$, matrix $\Theta\Sigma(\hat{\boldsymbol{n}})\Theta^T$ was invertable in every simulation and, as we can see in Figure B2, simulated $\Xi$ values are approximately chi-square distributed with 6 degrees of freedom.

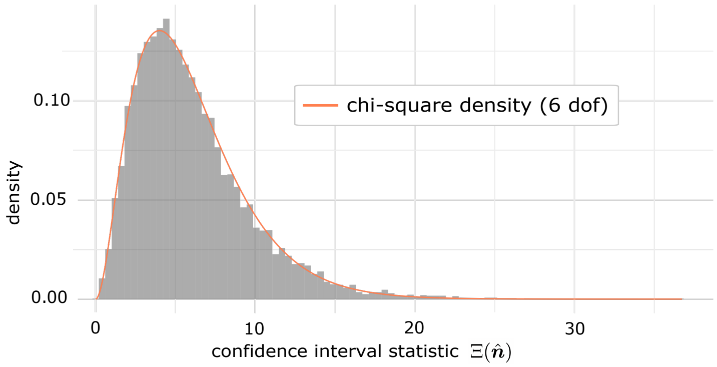


**Fig. B2** 10,000 simulated $\Xi$ values for simulation setting $s_A = s_B = s_C = 0.95$ and $n_{ABC} = 500, n_A = n_B = s_C = 50$ (see Section B.2) approximately follow a chi-square distribution with 6 degrees of freedom.

| $s_A = s_B = s_C$ | $n_{ABC}$ | $n_{AB} = n_{BC} = n_A = n_B = n_C$ | Empirical Coverage |
|---|---|---|---|
| 0.80 | 50 | 50 | 0.8879 |
| 0.80 | 100 | 50 | 0.8884 |
| 0.80 | 500 | 50 | 0.8955 |
| 0.85 | 50 | 50 | 0.8893 |
| 0.85 | 100 | 50 | 0.8919 |
| 0.85 | 500 | 50 | 0.8897 |
| 0.90 | 50 | 50 | 0.8906 |
| 0.90 | 100 | 50 | 0.8984 |
| 0.90 | 500 | 50 | 0.8963 |
| 0.95 | 50 | 50 | 0.8898 |
| 0.95 | 100 | 50 | 0.8953 |
| 0.95 | 500 | 50 | 0.8940 |

**Table B1** Empirical coverage for the simulation of different triplet, pair and singlet abundances $\boldsymbol{n}$ and labeling efficiencies $s_A, s_B, s_C$ (see Section B.2) at a theoretical coverage of $1 - \alpha = 0.90$.

### C ConditionalColoc Comments

We experienced that ConditionalColoc, although aiming to output probabilities, in some cases yields values greater than one and hence the errors in relative abundance detection are not bounded by one as well. In the following Figure C3 we show the ConditionalColoc outliers not depicted in Figure 2.

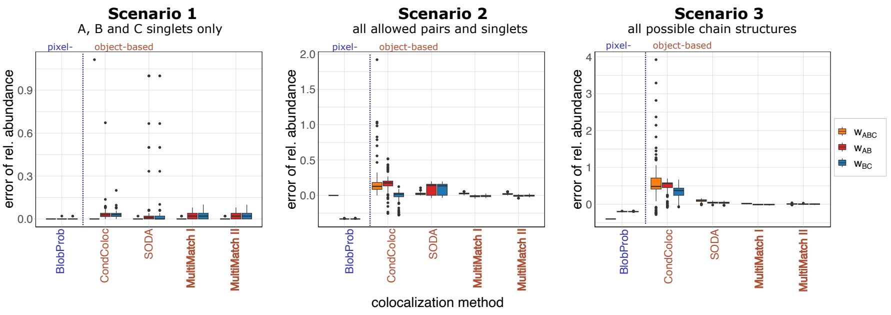


**Fig. C3** **Simulation study with ConditionalColoc outliers.** As Figure 2 but including outliers of ConditionalColoc resulting in errors in relative abundances greater than one. In each Scenario 100 STED images and different abundances of triplets, pairs and singlets were simulated with 100% labeling efficiency. **A.** Method specific boxplots of the errors in detected relative (scaled by the total number of points in channel B) structure abundances are displayed. The error is computed by subtracting true relative abundance from detected relative abundances. In *Scenario 1* only A,B and C singlets, in *Scenario 2* all possible singlets as well as AB and BC pairs and in *Scenario 3* ABC triplets, AB, BC pairs and A, B and C singlets were simulated.

### D MultiMatch Mode II: Experimental Settings

To compare MultiMatch Mode I and II, we repeated the colocalization analysis of experimental STED image settings with MultiMatch Mode II (see Figure D4). As expected, the mean relative abundance curves $\hat{\mathbf{w}}$ and the corresponding estimation results of $\hat{\mathbf{n}}$ are comparable to the results reported by Mode I (see Figure 4 for analysis output by Mode I). However, since Mode II does not prioritize the detection of triplets, unlike Mode I, the triplets frequencies are more underestimated in Setting 3, where only triplets should occur in the image. The maximal relative mean abundance of detected ABC triplets, which is attained for colocalization threshold $t = 10$ pixels is $w_{ABD} = 0.77$. For MultiMatch Mode I it is closer to the (by experimental design known) truth of having triplets only, by reaching a maximal relative abundance of $w_{ABC} = 0.8$ (see Figure 4).

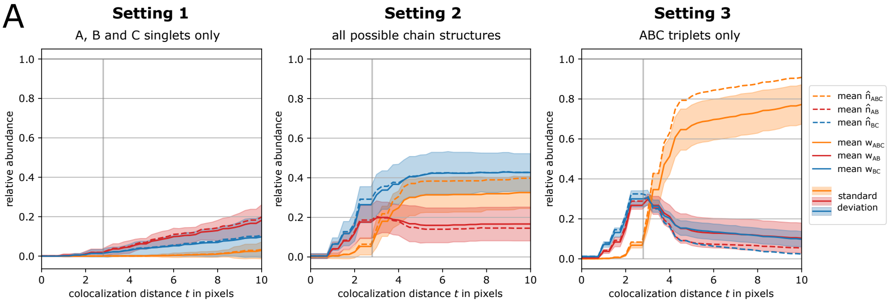


**Fig. D4 MultiMatch Mode II relative abundance curves $\boldsymbol{w}(t)$ for experimental STED images.** As Figure 4 but analysed with MultiMatch Mode II instead of Mode I. For each setting the solid curves are mean relative abundances with standard deviation bands across a range of colocalization thresholds $t$ from 0 to 10 pixels (25 nm = 1 pixel). The abundances are scaled by the total number of points detected in channel B. Additionally, incomplete labeling efficiency (90% in each channel) corrected abundances are plotted as dotted curves. The true colocalization threshold of 70 nm within nanoruler structures is depicted as a vertical line. **A.** *Setting 1* contains singlet, *Setting 2* triplet, pair and singlet and *Setting 3* triplet nanorulers only.

### E Additional Analysis: Simulated Scenarios

#### E.1 Method comparison across colocalization thresholds

We also tested the performances of considered colocalization methods across different colocalization thresholds $t$ and show results in Figure E5. Methods were evaluated on the colocalization grid $t \in \{1, 2, 3, 4, 5, 6, 7, 8\}$ pixels. We experienced, that BlobProb and ConditionalColoc were not directly applicable to a batch of images at once. In particular, BlobProb requires the user to load every image separately into an ImageJ/Fiji Graphical User Interface, where parameters as the colocalization threshold have to be adjusted by hand, respectively. For the MATLAB implementation of ConditionalColoc, all images within a simulation scenario had to be combined into a 'movieList', which could then be input as a whole for colocalization analysis. However, the analysis had to be performed separately for each colocalization threshold. With the runtime of 2 minutes per image and colocalization radius, as reported in the ConditionalColoc manual (https://github.com/kjaqaman/conditionalColoc), the evaluation of our simulation study with ConditionalColoc took about 1000 times longer than with our MultiMatch implementations (0.1 seconds per image and colocalization radius, see Methods Section 4.2).

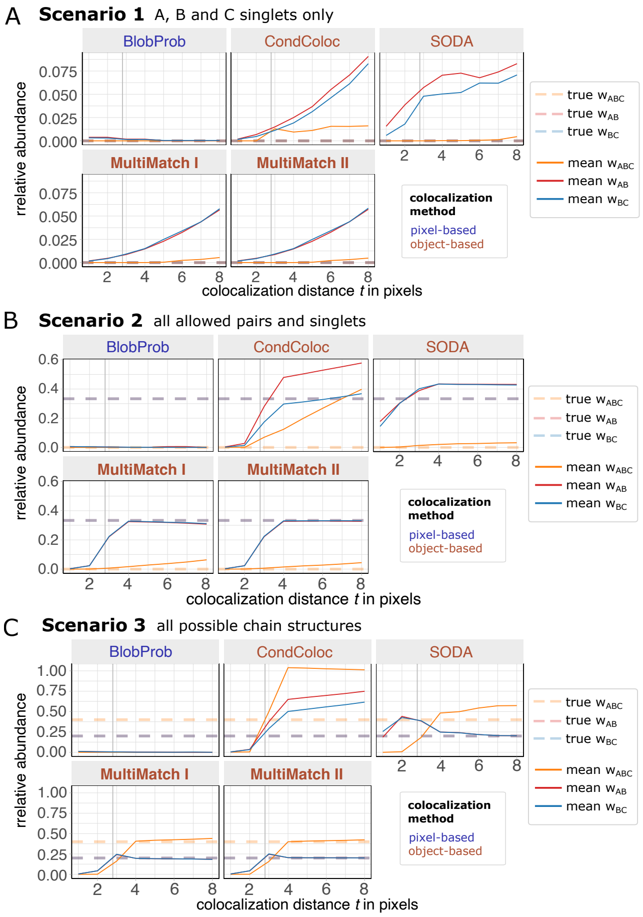


**Fig. E5 Simulation study for three combinations of chain structures along different colocalization thresholds *t*.** Simulation setup described in Section 2.2: In each Scenario 100 STED images and different abundances of triplets, pairs and singlets were simulated with 100% labeling efficiency. Mean relative abundances curves (scaled by the total number of points in channel B) are shown per colocalization method and chain structure. True simulated relative abundances are plotted as transparent, horizontal dashed lines. **A.** In *Scenario 1* only A,B and C singlets, in **B.** *Scenario 2* all possible singlets as well as AB and BC pairs and in **C.** *Scenario 3* ABC triplets, AB, BC pairs and A, B and C singlets were simulated.

#### E.2 Comparison with Nearest Neighbor Matching

In addition to the considered colocalization methods provided as packages, plugins or executable scripts, one can also consider greedy Nearest Neighbor Matchings as a compatible algorithm to MultiMatch.

Nearest Neighbor Matchings can be implemented in several ways depending on the order of points within a channel and the order of channels being matched in pairs. For the method comparison below, we used the following implementation:

1. For each point in channel A, assign it to its nearest neighbor in B as soon as their pairwise distance is smaller than the colocalization threshold $t$. If matched, do not consider the respective B point for further nearest neighbor searches of channel A points. The match is stored as **AB pair**.
2. Repeat 1. to match points in channel B to their nearest neighbors in channel C. Respective matches are stored as **BC pairs**.
3. If an AB and a BC pair share the same B point, they are re-annotated into one **ABC triplet**.

We applied MultiMatch Mode I and II and the above described Nearest Neighbor Matching approach directly to the simulated point clouds without additional conversion to microscopy intensity images. To illustrate the differences between MultiMatch's global optimization procedure and the effect of local, greedy Nearest Neighbor searches, we chose to simulate settings with a high particle density and an especially high number of triplets:

**Scenario i:** 1000 ABC triplets only.
**Scenario ii:** 1000 ABC triplets and 1000 B singlets.
**Scenario iii:** 1000 ABC triplets, 500 A, B and C singlets, respectively.

As in all other simulation scenarios, the pixel size was simulated as 1 pixel = 25 nm, the image size was set to 400×400 pixels and the true colocalization distance was fixed to $t$ = 70 nm. For each setting 100 images were simulated, but we directly evaluate the coordinates of simulated point clouds without further translation into an intensity image nor simulation of microscopy noise or point spread function convolution (see examples in Figure E6D).

As can be seen in Figure E6A, in Scenario i the Nearest Neighbor Matching approach underestimates the number of ABC triplets and overestimates the abundance of AB and BC pairs, although only triplets were simulated. For the a colocalization threshold of 4 pixels = 100 nm, the mean abundance detected by the Nearest Neighbor Matchings only reaches around 939.43 of 1000 simulated ABC triplets with a standard deviation of ca 0.197, while in both MultiMatch Modes I and II all 1000 simulated ABC triplets are recovered for every simulated image.

In Scenario ii we can showcase a similar behavior: Nearest Neighbor Matchings only reach a maximal average abundance of 838.71 out of the 1000 simulated ABC triplets for $t$ = 4 pixels.

Additionally, we can observe that this setting is also challenging MultiMatch Mode II: Similar to Nearest Neighbor Matchings, MultiMatch Mode II underestimates triplets due to the disproportionate abundance of Type B particles and the dense particle distribution. Mode II finds more AB pairs with lower overall pairwise particle distances and therefore on average misses around 15% of all simulated ABC triplets in maximal colocalization threshold.

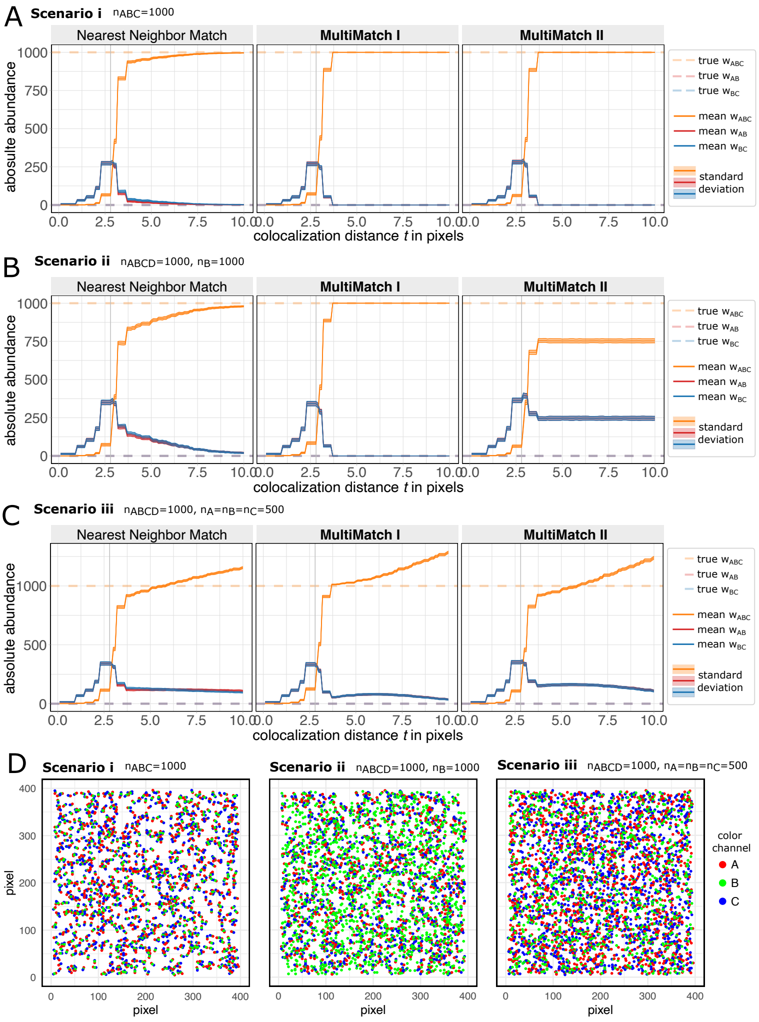


**Fig. E6 Simulation study for three combinations of chain structures along different colocalization thresholds** $t$**.** The simulation setup is described in detail in Section E.2. Mean absolute abundances curves with standard deviation bands are shown per method and chain structure. True simulated relative abundances are plotted as transparent, horizontal dashed lines. **A.** In *Scenario i* only ABC triplets and in **B.** *Scenario ii* ABC triplets as well as B singlets were simulated. **C.** *Scenario ii* ABC triplets as well as A,B and C singlets were simulated. **D.** Representative particle clouds for Scenarios i-iii.

Interestingly, the shape of abundance curves for the Nearest Neighbor Matching is very similar between Scenarios i, ii and iii altough in the first we only simulated triplets and in the latter two we added different types and ratios of singlets. The Nearest Neighbor Matching approach does not only underestimate triplet abundances but also shows no clear plateau to discriminate between different colocalization structures. MultiMatch Mode I abundance curves, on the other hand, stabilizes at the correct abundance for triplet abundances for Scenario i and ii. If a random distribution of more than one type of singlets is present in the image (Scenario iii), singlets are matched as soon as the colocalization threshold is high enough. Still, one can observe that the abundance curve slopes visibly drop for $t \geq 4$ pixels.

#### E.3 Additional Four-Color STED Simulation Scenarios

Supporting Section 2.5, we additionally simulated the following three four-color STED image scenarios:

**Scenario III:** 50 A, B, C and D singlets, respectively, and no further quadruples, triplets nor pairs.
**Scenario IV:** 200 ABCD quadruples and 100 A,B, C and D singlets, respectively.
**Scenario IV:** 500 ABCD quadruples only.

From the respective MultiMatch Mode II results of those two simulations scenarios, shown in Figure E7, one draws that abundance curves again are stabilizing after a colocalization threshold of $t = 4$ pixels. In case the detection of chain structures, here ABCD quadruples, is aggravated by incomplete labeling efficiency, our estimation framework leads to a consistent improvement of detection results towards the simulated ground truth.

For simulation Scenarios IV and V it becomes obvious, that due to high number of particles in the image, the simulated noise and according point detection errors, MultiMatch can not recover all simulated quadtrupels. In this case, the interactive napari viewer (napari contributers, 2019) can help to evaluate the noise level, point detection performance and matching results. For uniform noise levels and point detection errors across all channels, we recommend to evaluate channel-wise scaled instead of absolute abundance curves.

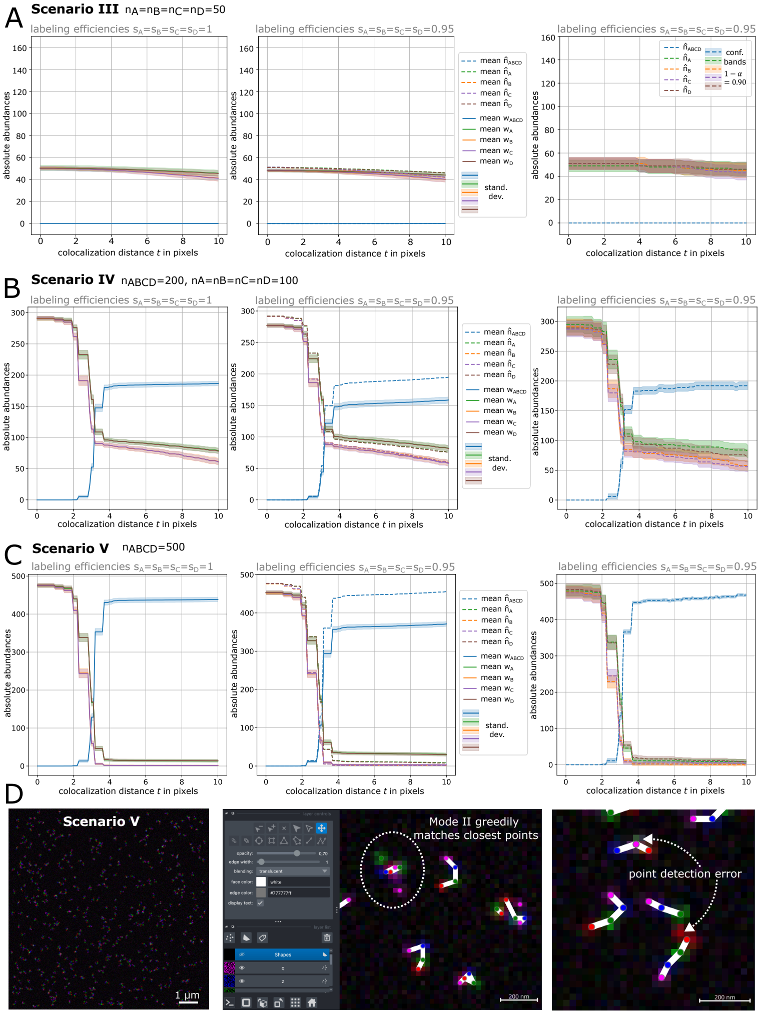


**Fig. E7 MultiMatch Mode II abundance curves $w(t)$ and estimation results $\hat{\boldsymbol{n}}(t)$ for simulated four-colour STED images.** For each scenario, images with complete labelling efficiency (left) and with incomplete labelling efficiency (middle) were simulated. Solid curves are mean absolute detected abundances with a standard deviation bands. Corrected abundances are plotted as dotted curves. For one exemplary STED image simulated with incomplete labelling efficiency, corrected abundance curves and corresponding confidence bands are shown (right). **A.** *Scenario III:* Only singlets were simulated. **B.** *Scenario IV:* ABCD quadruplets and A,B,C and D singlets were simulated. **C.** *Scenario IV:* ABCD quadruplets only were simulated. **D.** Representative STED image for Scenario V with image details in the interactive napari viewer allowing a visual check of the image and MultiMatch output quality.
